# Clinical and molecular characterization of an outbreak of leptospirosis in dogs from Los Angeles County, California, USA, 2021

**DOI:** 10.64898/2026.03.24.706307

**Authors:** Max W. Randolph, Jarlath E. Nally, Sean K. Yoshimoto, Betty Chow, David M. Wagner, Nathan E. Stone, Jason W. Sahl, Camila Hamond, Karen LeCount, Tod Stuber, Hans van der Linden, Krystle Reagan, Alexander Schrieber, Jamie Sebastian, Jane E. Sykes

## Abstract

**Background:** In 2021, the Los Angeles County (LAC) Department of Public Health suspected a leptospirosis outbreak in LAC affecting over 200 client-owned dogs.

**Objective:** To characterize the outbreak and describe microbiologic findings, risk factors, diagnostic test performance, and outcomes in dogs diagnosed with leptospirosis at two specialty practices.

**Methods:** *Leptospira* culture isolates from four cases were subjected to serotyping and whole genome sequencing (WGS); WGS was also performed on one enriched genome isolate. After the outbreak, data were gathered on 59 cases through record review and compared to the background hospital population (controls, n=15,536).

**Results:** All isolates were *Leptospira interrogans* serovar Canicola, but each was distinct based on WGS. Cases clustered in space and in time. Cases evaluated during the outbreak peak had increased odds of exposure to indoor congregate facilities (ICFs). None of 47 dogs with known leptospirosis vaccination history were completely vaccinated. *Leptospira* real-time PCR on blood and urine and initial serologic testing using the microscopic agglutination test and point-of-care tests were positive in 15/56 (27%), 49/54 (91%) dogs, 22/29 (76%), and 27/35 (77%) dogs respectively. Fifty-four (92%) of 59 dogs survived to discharge; some remained azotemic. No associated human cases were identified.

**Conclusions and Clinical Importance:** *L. interrogans* serovar Canicola was associated with a leptospirosis outbreak in unvaccinated dogs from LAC, which had public health implications given widespread dog ownership rates. Data analysis suggested multiple infection sources, including ICFs. Urine PCR was the most sensitive diagnostic test. Such outbreaks might be prevented through more widespread vaccination.

## INTRODUCTION

Leptospirosis is a multisystemic zoonotic disease of humans, domestic dogs, and other mammalian host species caused by pathogenic spirochete bacteria of the genus *Leptospira*, which are chronically shed in urine of a variety of subclinically-infected reservoir hosts, especially rodents.^1–3^ Several different serovars of *Leptospira interrogans* and *Leptospira kirschneri* have been identified in dogs with leptospirosis, including serovar Canicola,^4–7^ for which domestic dogs have also been considered the primary reservoir host. Clinically, severe leptospirosis in dogs resembles Weil’s disease in humans; signs include fever, lethargy, inappetence, polydipsia, polyuria, vomiting, diarrhea, and icterus due to acute kidney injury (AKI) and cholestatic hepatopathy.^1^

Worldwide, leptospirosis is more prevalent in subtropical regions with heavy rainfall. In the United States, seasonality in dogs is associated with regional precipitation patterns.^1,8,9^ In both humans and dogs, most infections occur when spirochetes in contaminated water or soil penetrate intact mucous membranes or abraded skin. Less often, direct transmission can follow exposure to contaminated urine from reservoir hosts, bite wounds, venereal transmission, or predation.^1,10–12^ Most cases are sporadic, but outbreaks in humans often follow flooding or sporting events involving water.^13^ Transmission between incidental hosts is rarely described, perhaps because of reduced shedding compared to reservoir hosts.^14^ For almost a century, dogs have been implicated as an important reservoir for transmission of *L. interrogans* serovar Canicola to humans, especially in resource-poor countries where free-roaming dogs are widespread,^1,15–18^ although molecular proof of transmission is lacking and evidence for widespread subclinical carriage has predominantly been based on serologic data, which does not reliably predict the infecting serovar.^19^ Further, chronic and subclinical shedding of serovar Canicola has also been identified in other species, including rodents, pigs, and cattle.^16,20–22^ Before 2023, vaccination of dogs for leptospirosis was reserved for dogs considered at-risk based on geographic region and lifestyle.^1^ Current vaccines for dogs are serovar-specific bacterins; in North America these include serovars Canicola, Icterohaemorrhagiae, Grippotyphosa, and Pomona.

Diagnosis of leptospirosis is optimized by combining serologic tests and pathogen-detection tests (real-time PCR and culture).^23^ Acute and convalescent phase serologic testing using the microscopic agglutination test (MAT) is widely accepted as the gold standard test in humans and dogs but is laborious to perform, often negative in acutely ill patients, and requires maintenance of a large panel of live pathogenic *Leptospira* serogroups to optimize sensitivity.^23^ Culture requires special media, expertise, and often weeks or months of incubation, so is not widely used for routine diagnosis; however, it is required for accurate serovar identification. Because spirochetes are found in the blood during the first week of illness, and later appear in urine, testing both blood and urine using PCR optimizes clinical sensitivity,^23^ although the relative clinical sensitivity of blood versus urine PCR is poorly understood. In North America, point-of-care lateral-flow chromatographic assays are commercially available for detection of antibodies to pathogenic leptospires in dogs (e.g., IgM [WITNESS Lepto Rapid Test, Zoetis, Parsippany, NJ], IgG and IgM [SNAP Lepto, IDEXX, Portland, ME]).^1^ To date, clinical validation of these assays has been limited, and assay performance might vary regionally depending on circulating serovars.

In Los Angeles (LA) County, California, laboratories and veterinarians must report leptospirosis to local public health authorities.^24^ In 2021, the LA County Department of Public Health (DPH) identified an outbreak of leptospirosis involving at least 201 client-owned dogs in west LA.^24^ Many dogs developed illness within days of being boarded in one of two regional dog daycare facilities from the region, although specific details were not available. Because affected dogs often seroconverted with very high MAT titers to serogroup Canicola, a serovar belonging to serogroup Canicola was suspected as the cause. It was therefore proposed that the outbreak was fueled by dog-to-dog transmission via direct contact with infected urine.^25^ Within a 6-month period, more than 50 affected dogs were examined at two major veterinary specialty hospitals in the region, where extensive diagnostic workups and treatment occurred. We sought to use this unique opportunity to accurately identify the causative agent and characterize risk factors, and describe clinical findings, diagnostic test results, and outcomes.

## MATERIALS AND METHODS

### *Leptospira* Isolation and Identification

At the time of the outbreak (September 2021), we contacted veterinarians at two veterinary emergency and specialty practices in west LA, located 0.6 km apart: VCA Animal Specialty and Emergency Center [ASEC] and VCA West LA Animal Hospital [WLA]. We requested blood and urine specimens from dogs that were highly suspected to have leptospirosis for concurrent *Leptospira* culture, real-time PCR, and serologic testing. Dogs were excluded from this part of the study if antimicrobial drug therapy had commenced. In preparation for specimen collection, 5-mL aliquots of *Leptospira* Hornsby-Alt-Nally (HAN)^26^ liquid media, HAN semi-solid media, and T80/40/LH semi-solid media^26^ were shipped to WLA overnight on ice. Uninoculated media was stored at 4°C. Specimens were collected by cystocentesis or direct venipuncture. In October and November 2021, the semi-solid media were inoculated at point-of-care with 2-3 drops of either urine or blood from four dogs (LAD1 and LAD3-LAD5). Liquid HAN media was inoculated with 1 mL urine but not blood. Inoculated media, blood, and urine were immediately shipped at ambient temperature to the National Centers for Animal Health (NCAH) (United States Department of Agriculture, Ames, IA). There, aliquots of media were subjected to direct fluorescent antibody (DFA) examination and *Leptospira* culture using HAN liquid (37°C), HAN semisolid (37°C), and T80/40/LH semisolid media (29°C). Blood and urine were subjected to real-time *lipL32* gene PCR.^27,28^ Inoculated tubes were examined daily by darkfield microscopy. Isolates were subjected to whole genome Illumina sequencing and serotyping was performed using microscopic agglutination as previously described.^29^ Serovar typing of strains LAD1, LAD3, LAD4 and LAD5 was performed at the WOAH Reference Laboratory for Leptospirosis, University of Amsterdam, The Netherlands, by the MAT with panels of monoclonal antibodies (mAbs) that characteristically agglutinate serovars from the serogroup Canicola, as previously described.^30^

### *Leptospira* Genome Enrichment

#### DNA capture and enrichment methods

For one dog (LAD2) inoculated media was not provided, but *lipL32* gene real-time PCR was positive on urine (Ct = 26.3). To obtain a genome from this sample, we performed *Leptospira* genome capture and enrichment, as previously described.^31,32^ Extracted DNA was diluted to ∼4 ng/µL in 10mM Tris-HCL to a final volume of 40 µL and sonicated to an average size of 162 bp using a Q800R2 sonicator (QSonica, Newtown, CT, US). A dual indexed library was then prepared using Agilent Sure-Select methodology (XT-HS kit, Agilent, Santa Clara, CA, US). One round of DNA capture and enrichment was performed. The enriched library was sequenced on an Illumina MiSeq instrument using a MiSeq v3 600 cycle kit (2×300 base pair reads).

#### Read classifications

To estimate the percentage of *Leptospira* reads in the enriched sequences, reads were mapped against the standard Kraken database with Kraken v2.1.2.^33^

#### Genome assemblies, read mapping, and phylogenomics

Sequencing reads were assembled for LAD1 and LAD3-LAD5 using SPAdes v3.13.0^34^ with default settings, and multi-locus sequence type (ST) was determined by querying the assemblies against the pubMSLT (scheme 1) database.^35^ Two phylogenies were created. First, single nucleotide polymorphisms (SNPs) were identified among the four isolate genomes (LAD1, LAD3, LAD4, and LAD5), one enriched genome (LAD2), and 13 publicly available *L. interrogans* serovar Canicola genomes (GenBank accession numbers provided in **Figure 1**) by aligning LAD2 enriched reads against the LAD1 assembly using minimap2 v2.22.^36^ SNPs were called from the BAM file with GATK v4.2.2^37^ using a depth of coverage ≥10x and a read proportion of 0.9. SNPs that fell within duplicated regions, based on a reference self-alignment with NUCmer v3.1,^38^ were filtered from downstream analyses. All methods were wrapped by NASP v1.2.1.^39^ Maximum likelihood phylogenies were then inferred on the concatenated SNP alignments using IQ-TREE v2.2.0.3 with default parameters, 1,000 bootstraps replicates,^40^ and the integrated ModelFinder method.^41^ The phylogeny was rooted with *L. interrogans* serovar Pomona strain Kennewicki LC82_25 (GCA_000243635.3). To determine breadth of coverage, enriched reads were aligned against reference genome *L. interrogans* serovar Pomona strain Kennewicki LC82_25 with minimap2, and the per-base depth of coverage was calculated with Samtools v1.6.^42^ Second, to investigate fine-scale differences among the five dog genomes (LAD1-5), a maximum likelihood phylogeny was created as described above except using the “-fast” option in IQ-TREE and only LAD1-5 genomes were included (LAD1 was used as the reference). This phylogeny was midpoint rooted.

**Figure 1.**
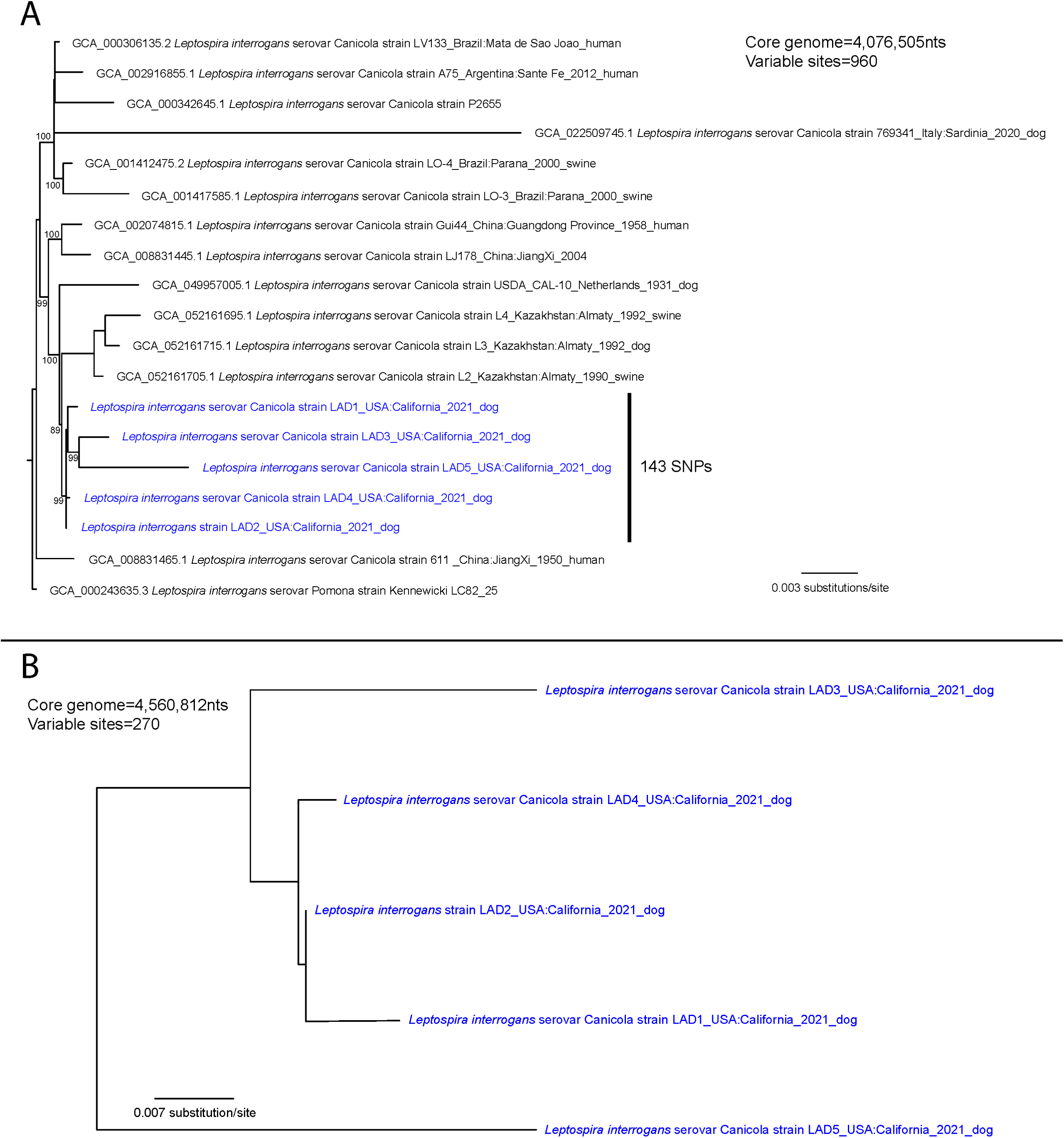
*Leptospira interrogans* serovar Canicola whole genome phylogenies. A) Whole genome maximum likelihood phylogeny of four *Leptospira interrogans* serovar Canicola isolate genomes and one enriched genome from dogs (blue text) with 13 publicly available *Leptospira interrogans* serovar Canicola genomes based upon a concatenated SNP alignment of 960 variable positions within a shared core genome of 4,076,505 nucleotides (nts). The phylogeny was rooted with reference genome *L. interrogans* serovar Pomona strain Kennewicki LC82_25. The canine isolates group together into a unique clade defined by three novel SNPs and displaying 99% bootstrap support (bootstrap values indicated on branch nodes). B) Whole genome maximum likelihood phylogeny based upon an alignment of 270 variable nucleotide positions within a shared core genome of 4,560,812nts using five dog genomes generated from isolates and via DNA capture and enrichment. The phylogeny was midpoint rooted and illustrates the full SNP diversity within the core genome of these infecting strains.

#### Data Availability

Sequencing data and genome assemblies are available at NCBI under BioProject ID PRJNA1377681; accession numbers are listed in **Table S1**.

### Medical Records Review

In 2022, after the outbreak concluded based on reporting data gathered by the LA County DPH,^24^ we performed a search of electronic medical records from ASEC and WLA for client-owned dogs diagnosed with leptospirosis from April 2021 to December 2021 (inclusive). Each of the search terms “lepto*,” “microscopic agglutination test,” “MAT,” and “lepto* PCR” were used. To be included in the study, dogs had to have two or more of the following clinical signs consistent with leptospirosis: lethargy, polyuria or polydipsia, hyporexia or anorexia, vomiting, or diarrhea. In addition, diagnosis of leptospirosis required detection of *Leptospira* DNA in blood or urine using PCR, a single MAT titer ≥ 1:6400, or a 4-fold or greater rise in MAT titers. Real-time PCR was performed at either IDEXX Laboratories (North Grafton, MA, USA)(ASEC) or Antech Diagnostics (WLA). Serologic testing using MAT was performed at either IDEXX Laboratories (ASEC) or Michigan State University (through Antech Diagnostics) (WLA) using their standard laboratory testing protocol for diagnostic specimens covering six serovars: Autumnalis, Bratislava, Canicola, Grippotyphosa, Icterohaemorrhagiae, and Pomona. Both laboratories participate in the International Leptospirosis Proficiency Testing Scheme.^43^

Information extracted from medical records included *Leptospira* vaccination history; zip code of residence, date of initial examination and duration of illness; age (years), sex, breed, and body weight (kg); clinical signs; the results of hematology, serum biochemistry, and urinalysis (hematocrit; neutrophil count; platelet count; serum concentrations of urea nitrogen, creatinine, and total bilirubin; serum activities of ALT and ALP; presence or absence of glucosuria); results of *Leptospira* real-time PCR and serologic testing using MAT, WITNESS Lepto (Zoetis), and SNAP Lepto (IDEXX Laboratories); duration of hospitalization; antimicrobials administered for treatment of leptospirosis; and outcome (survival, death or euthanasia). Based on frequency of case presentations at the two hospitals, case data were organized into three periods relative to the peak of the epidemic curve (pre: 4/17/2021 – 6/4/2021, peak: 7/8/2021 – 9/8/2021, and post: 9/20/2021 – 12/5/2021).

#### Statistical Analysis

Univariate analysis was used to compare case variables (age, breed, sex, body weight) to those of the background hospital population (hereafter referred to as controls). Controls consisted of dogs examined at ASEC and WLA between April and December 2021 (inclusive). To protect confidentiality, data for controls were only available in categorical format. For continuous case data, normality was determined using a Shapiro-Wilk test. Categorical variables were compared using Fisher’s exact test. A Mann-Whitney U test was used to assess the relationship between serologic test results and duration of clinical signs. The Wilcoxon matched pairs signed rank test was used to compare initial serum creatinine concentration to serum creatinine concentration after treatment. All analyses were performed using GraphPad Prism v10.4.1; P-values ≤ 0.05 were considered significant.

Spatial analysis was performed using ArcGIS Pro 3.6.0 (Esri Inc., Redlands, CA). The spatial distribution of cases per 100,000 households was compared to that for controls at the zip code of residence level. Household data for zip code tabulation areas for 2021 were obtained from the United States Census Bureau. Zip code areas within a 15-20 km radius of the clinics (zip code 90025) were used for analysis, which included 57/58 (98.3%) of the leptospirosis cases from California and 15,536/23,731 (65.5%) of the hospital patient population. Global Moran’s I was used to assess for spatial autocorrelation, where a positive Z-score and significant P-values indicates spatial clustering. The Getis-Ord Gi* statistic (fixed distance band or inverse distance methods) was used to identify statistically significant spatial clusters of cold spots (low) and hot spots (high) across the entire outbreak as well by pre-peak, peak, and post-peak periods. Hot spots were compared for hospital patients and cases using a Hot Spot Analysis Comparison.

## RESULTS

### *Leptospira* Isolation and Identification

*Leptospira interrogans* serovar Canicola ST34 was isolated in media from four of the five dogs (LAD1, and LAD3-5) from which blood and urine were submitted for culture. All isolates were obtained from urine. The results of DFA, real-time PCR, and culture are shown in **Table 1**. Specimen collection dates, zip codes of residence, and exposure histories for the affected dogs are shown in **Table S2.** For the LAD2 urine specimen, after one round of enrichment, 99.4% (1,456,535/1,464,826) of sequencing reads assigned to *Leptospira* with a breadth and depth (minimum 10x) of coverage across the reference genome (*L. interrogans* serovar Pomona strain Kennewicki LC82_25) of 96.1% and 67.8x, respectively. Phylogenetic analyses revealed that leptospires in these five dogs (four genomes from cultures and one enriched genome) shared a common ancestry (as indicated by the shared presence of three novel SNPs), but were not identical, displaying 143 SNPs among them (**Figure 1A**). When comparing the core genome among the five isolates (4,560,812 nucleotides), we identified 12-224 SNPs between them (**Figure 1B**). LAD1 and LAD2 were the most similar (separated by 12 SNPs and both from the West Hollywood region).

**Table 1.** Results of direct fluorescent antibody, PCR, and culture in three different media types for *Leptospira* spp. on five dogs suspected to have leptospirosis during an outbreak of leptospirosis in Los Angeles County. Results are expressed as number positive/number of dogs tested.

| Specimen | DFA+ | PCR+ (Ct) | HAN liquid | HAN SS | T80/40/LH SS |
| --- | --- | --- | --- | --- | --- |
| Blood | 0/2 | 1/5 (38.9) | 0/3 | 0/3 | 0/3 |
| Urine | 2/3 | 5/5 (ND, 25, 25.7, 26.3, 31) | 3/5 (PID 2) | 4/5 (PID 2-12) | 4/5 (PID 2-19) |

DFA, direct fluorescent antibody; Ct, cycle threshold value; SS, semisolid; ND, not done; PID, post inoculation day when spirochetes identified by darkfield microscopy.

### Risk Factor

Fifty-nine dogs met the inclusion criteria: 23 from ASEC and 36 from WLA. When compared with the controls (15,536 dogs), cases had higher odds of being < 6 years of age (OR 8.41 [CI 3.91-18.09], *P* < .001), ≥ 15 kg (OR 4.35 [CI 2.44-7.76], *P* < .001), and male (OR 1.78 [CI 1.04-3.06], *P* = .04) (**Figures 2A-C**). Among female dogs, cases had higher odds of being intact compared to control dogs (OR 3.35 [CI 1.35-8.31], P = .02). There was no effect of intact status for males (OR 1.63 [CI 0.83-3.16], P = .18). Golden Retrievers (5/59, *P* < .02) and Siberian Huskies (3/59, *P* < .01) were overrepresented in the case population. Possible sources of exposure were identified in 45/59 (76%) dogs (**Table S3**). Thirty-one of 59 (53%) had exposure to ICFs; 24 (77%) of these 31 dogs were examined during the peak period. Time between most recent exposure to ICFs and onset of signs for 19 of the 30 dogs ranged from 2 to 35 days (median, 5 days). Cases in the peak period had higher odds of boarding or daycare than cases outside this window (OR 5.7 [CI 1.78 – 19.25], P < .01). Among cases, we found no association between sex and breed (Siberian Husky or Golden retriever) and exposure to ICFs, but dogs < 6 years of age (OR 10.00 [CI 1.09 – 120.9], P = .03) and dogs ≥ 15 kg were more likely to have exposure to ICFs than dogs < 15 kg (OR 5.52 [CI 1.46 – 18.24], P = .02).

**Figure 2.**
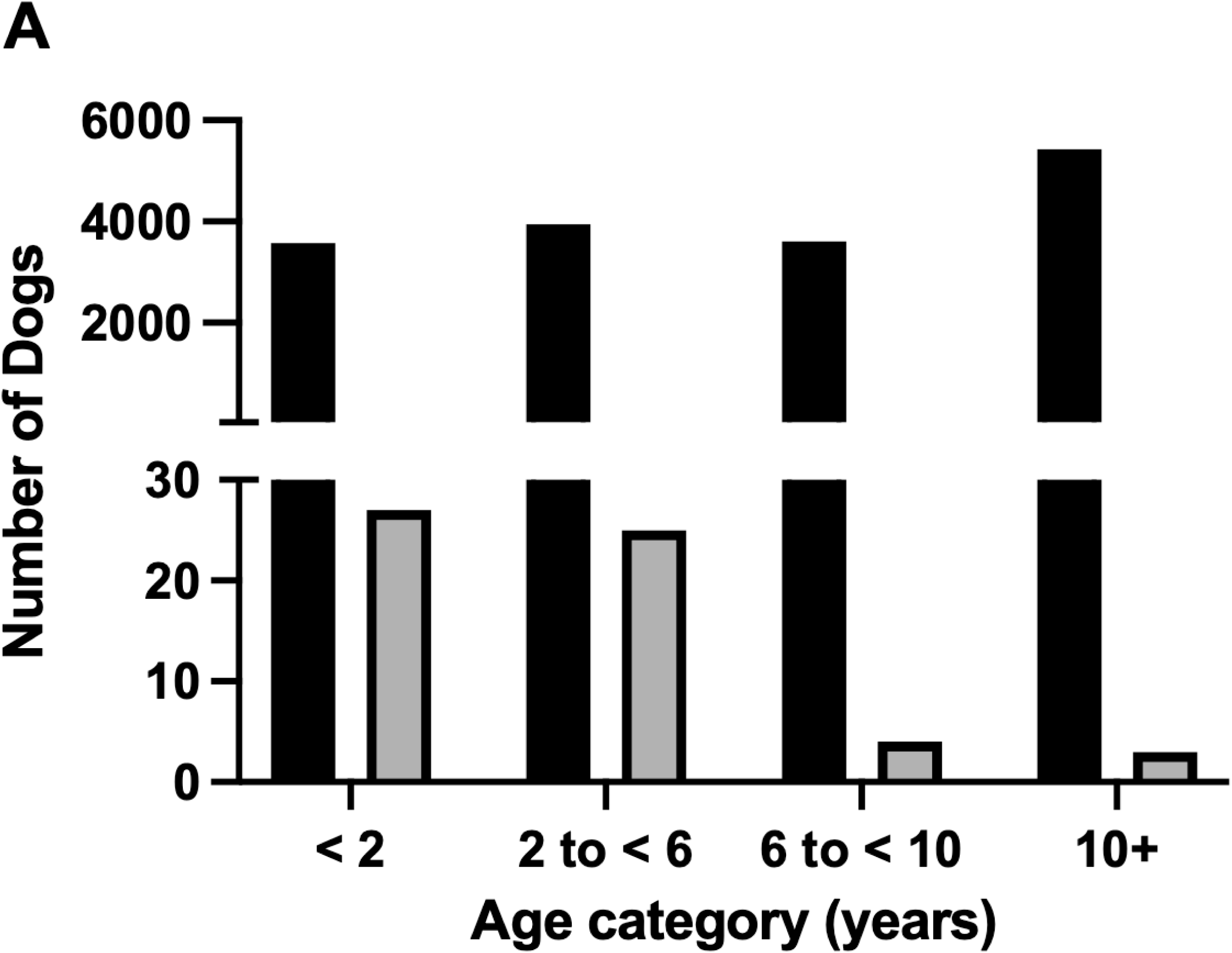
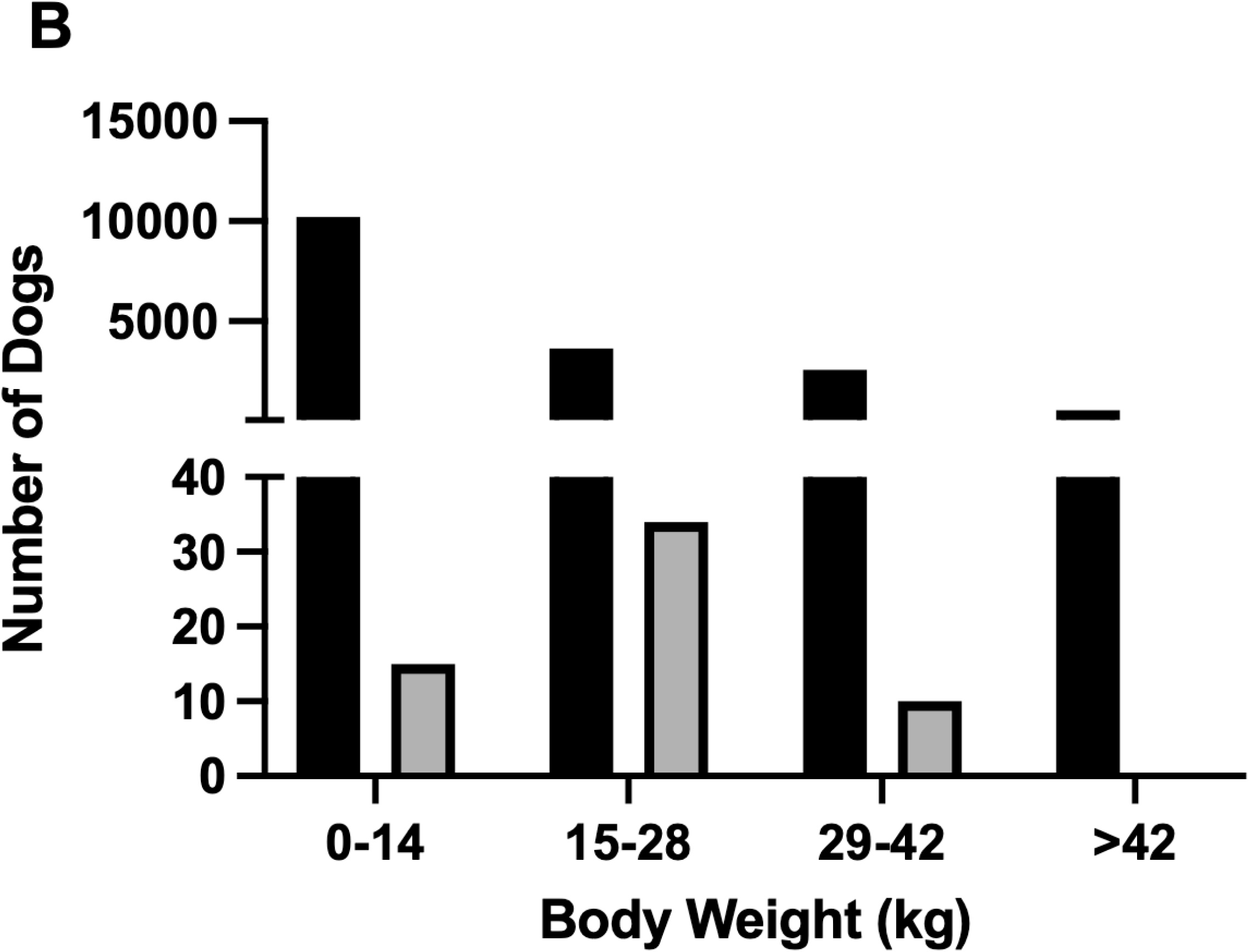
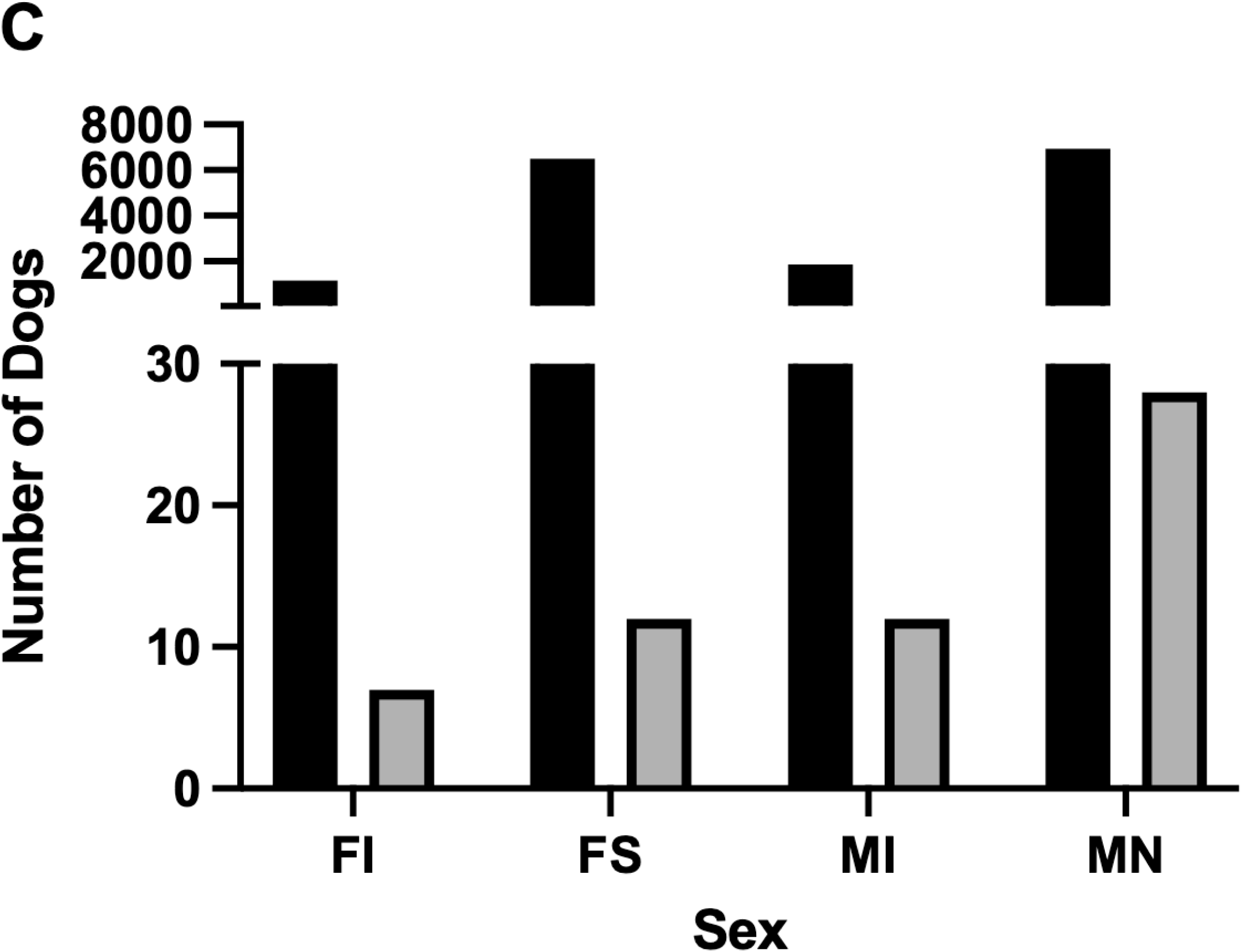
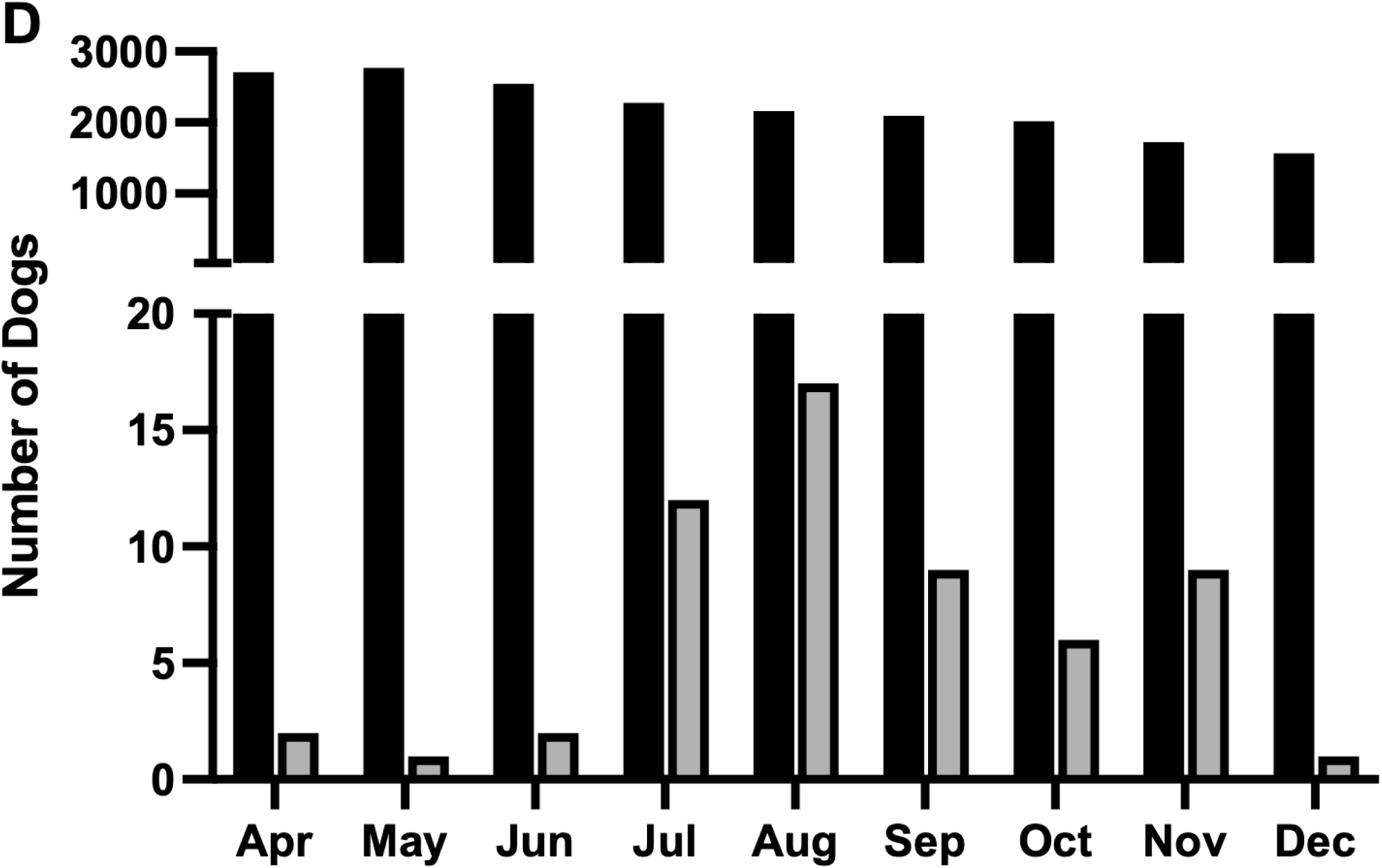
Age (**A**), body weight (**B**), sex (**C**), and month of evaluation (**D**) of 59 dogs with leptospirosis (grey bars) evaluated at 2 west Los Angeles specialty veterinary practices between April 1 and December 31, 2021, compared to those for the combined background hospital populations (black bars).

Of 47 dogs with a known *Leptospira* vaccination history, vaccines had never been administered to 41/47 (87.2%) dogs; 6/47 (12.8%) dogs had received a leptospirosis vaccine (all with Nobivac Lepto4, Merck Animal Health, NJ). Of the vaccinated dogs, four had only received the first of a 2-dose initial vaccination series before onset of illness and were not yet due for the second dose; one received the second dose 5 days before illness onset, and another four days after illness onset. The diagnosis for dogs that had received a vaccine was made using real-time PCR on urine (5) or blood (1); one of the dogs with a positive urine PCR had a positive MAT (1:25,600) nine days after its initial vaccine dose.

Thirty-four (58%) of cases were evaluated during the peak period. The temporal (**Figure 2D**) and spatial (**Figure 3**) distribution of cases differed from those of the control population, with cases clustering in zip code regions west and south of the clinics. We detected significant autocorrelation for cases (P = .01, Z-score = 2.5) and for controls (P < .001, Z-score = 5.6). Case hotspots shifted from regions concentrated around the clinics to regions southwest of the clinics during and after the peak of the epidemic, respectively (**Figure S1**).

**Figure 3.**
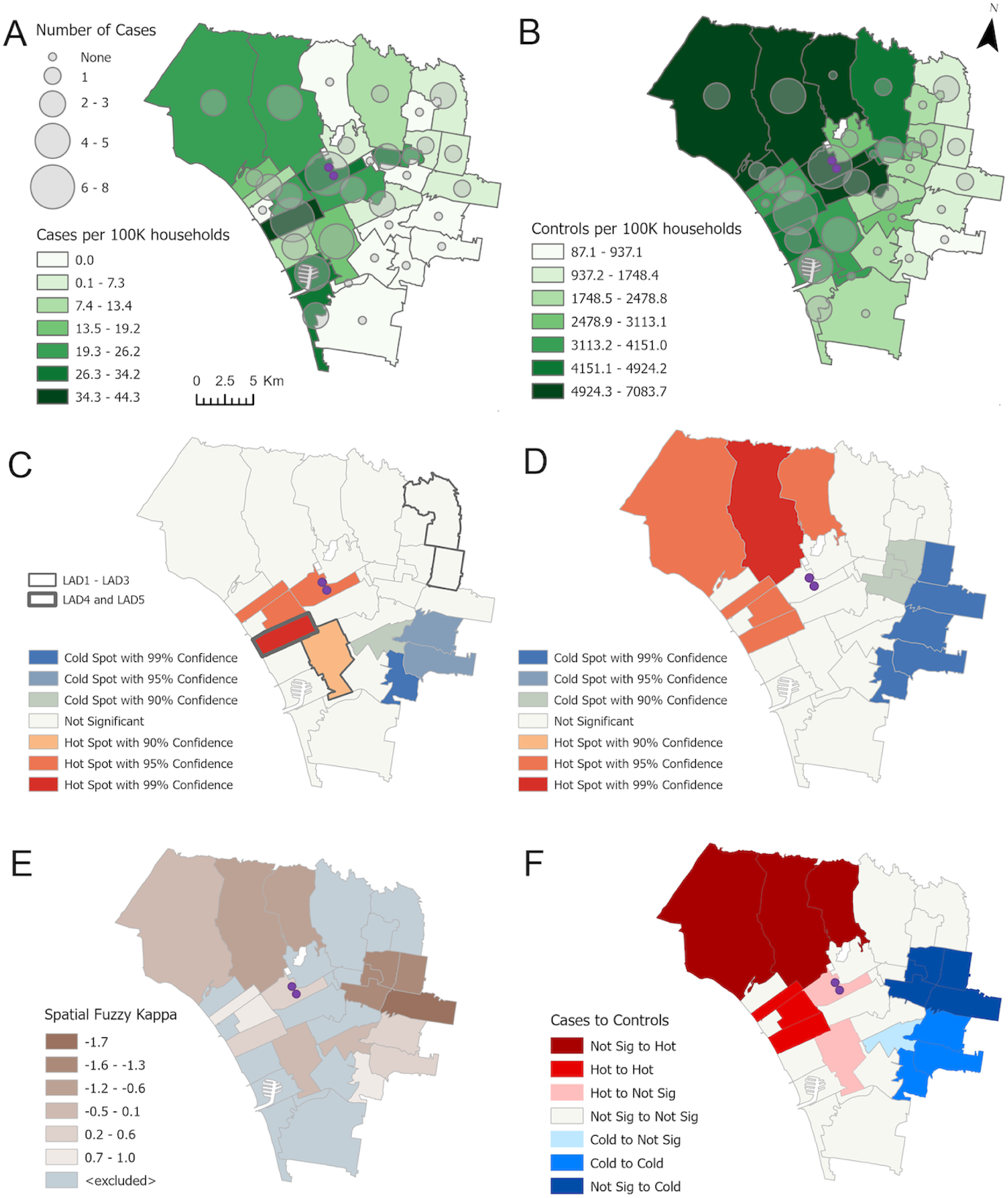
Maps of Los Angeles area used for spatial analysis (zip code regions within a 15 km radius of two veterinary specialty clinics [purple circles]). Graduated fill colors in zip code regions represent incidence data (cases and background hospital population [controls] per 100,000 households); transparent circles represent absolute case numbers. **A.** Leptospirosis cases in the study area (n = 57; two dogs fell outside the study area). **B.** Controls (n = 15,536). **C** and **D.** Distribution of hot spots and cold spots for case incidence and control incidence, respectively. In **C,** regions from which isolates were obtained are shown; the most similar isolates (LAD1 and LAD2) were from contiguous eastern zip codes. LAD4 and LAD5 were both from the same region in Santa Monica. **E** and **F.** Spatial fuzzy kappa (**E**) and significance comparisons (**F**) of hot and cold spots for case to control incidence. Spatial fuzzy kappa values of 1 indicate perfect association between hot and cold spots; and negative values indicate a negative relationship. The significance comparison compares significance categories between cases and the control population. Cold-to-hot, cold-to-nonsignificant, and hot to cold regions were not identified. Hot spots for cases are significantly distributed south and west of the clinic zip code region, whereas those for the hospital background population also include northern and eastern zip code regions. Sources: US Census Bureau; VCA, Inc.

### Clinicopathologic Findings

Clinical signs documented for > 10% of affected dogs were lethargy (45/59 [76%]), hyporexia (32/59 [54%]), vomiting (28/59 [47%]), polyuria/polydipsia (19/59 [32%]), anorexia (12/59 [20%]), and diarrhea (10/59 [17%]). Median duration of signs at initial examination was 3 days (range, 0-30 days). Laboratory abnormalities included anemia, neutrophilia, thrombocytopenia, azotemia, and biochemical evidence of cholestatic hepatic injury (**Table 2**). When results of both renal and hepatic function were available, 6/40 (15%) dogs had evidence of concurrent renal and liver function impairment and 29/40 (73%) had only evidence of renal function impairment; none had evidence of liver function impairment alone (**Table S4**). Urinalysis revealed glucosuria in 15/39 (38%) dogs. Seven dogs had thoracic radiographs; initial radiographs revealed no abnormal findings (n = 4) or mild diffuse interstitial to bronchointerstitial patterns (n = 3). For two dogs, follow-up radiographs were available; both had normal initial radiographs. One had a mild, diffuse interstitial pattern 2 days later; the other had a marked diffuse interstitial to alveolar pattern 7 days later that was consistent with leptospiral pulmonary hemorrhage syndrome.

**Table 2.**
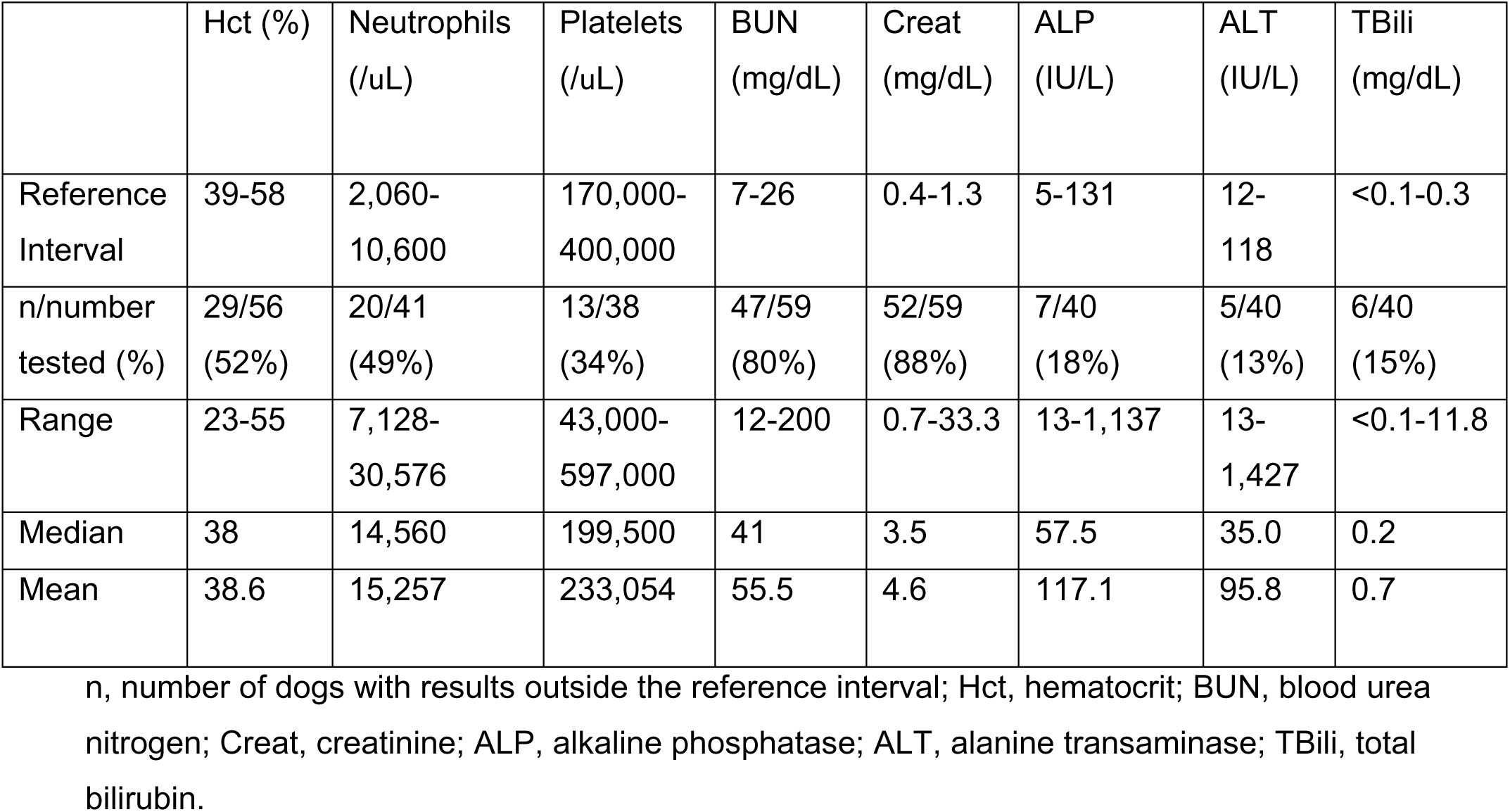
Hematologic and biochemical findings in 59 dogs diagnosed with leptospirosis during an outbreak in Los Angeles County.

### Nucleic Acid Amplification Testing

*Leptospira lipL32* gene real-time PCR was positive on whole blood in 15/56 (27%) dogs and urine in 49/54 (91%) dogs (OR 0.04, 95% CI 0.01 – 0.11, P < .001) (**Table 3**, **Figure 4**). In 52 dogs that had both blood and urine tested, nine (17%) had positive PCR results for both urine and blood.

**Table 3.** Median (range) days from illness onset to diagnostic test results in 59 dogs diagnosed with leptospirosis during an outbreak in Los Angeles County.

| Assay | n | Median (days) | Range (days) |
| --- | --- | --- | --- |
| Initial MAT |  |  |  |
| All | 29 | 5 | 0-30 |
| Highest $\geq 1:6400^*$ | 13 | 5 | 0-30 |
| Negative ($<1:50$) | 7 | 3 | 1-7 |
| Highest 1:100 to 1:3200 | 9 | 9 | 4-22 |
| Convalescent MAT $\geq 4$-fold increase or single $\geq 1:6400$ | 19 | 21 | 4-49 |
| WITNESS Lepto Rapid |  |  |  |
| All | 27 | 4 | 1-10 |
| Positive<sup>†</sup> | 20 | 4 | 1-9 |
| Negative | 7 | 3 | 2-10 |
| SNAP Lepto |  |  |  |
| All | 9 | 6 | 1-21 |
| Positive* | 8 | 6.5 | 3-21 |
| Negative | 1 | 1 | - |
| PCR |  |  |  |
| All | 58 | 4 | 0-21 |
| All blood | 56 | 4 | 0-21 |
| Positive blood | 15 | 3 | 1-14 |
| Negative blood | 41 | 4 | 0-21 |
| All urine | 54 | 4 | 0-21 |
| Positive urine | 49 | 4 | 0-21 |
| Negative urine | 5 | 4 | 2-21 |

MAT, microscopic agglutination test
\*One dog received the first dose of a leptospirosis vaccine 9 days before testing, but was also PCR-positive on urine
†One dog received the first dose of a leptospirosis vaccine 10 days before testing, but was also PCR-positive on urine

**Figure 4.**
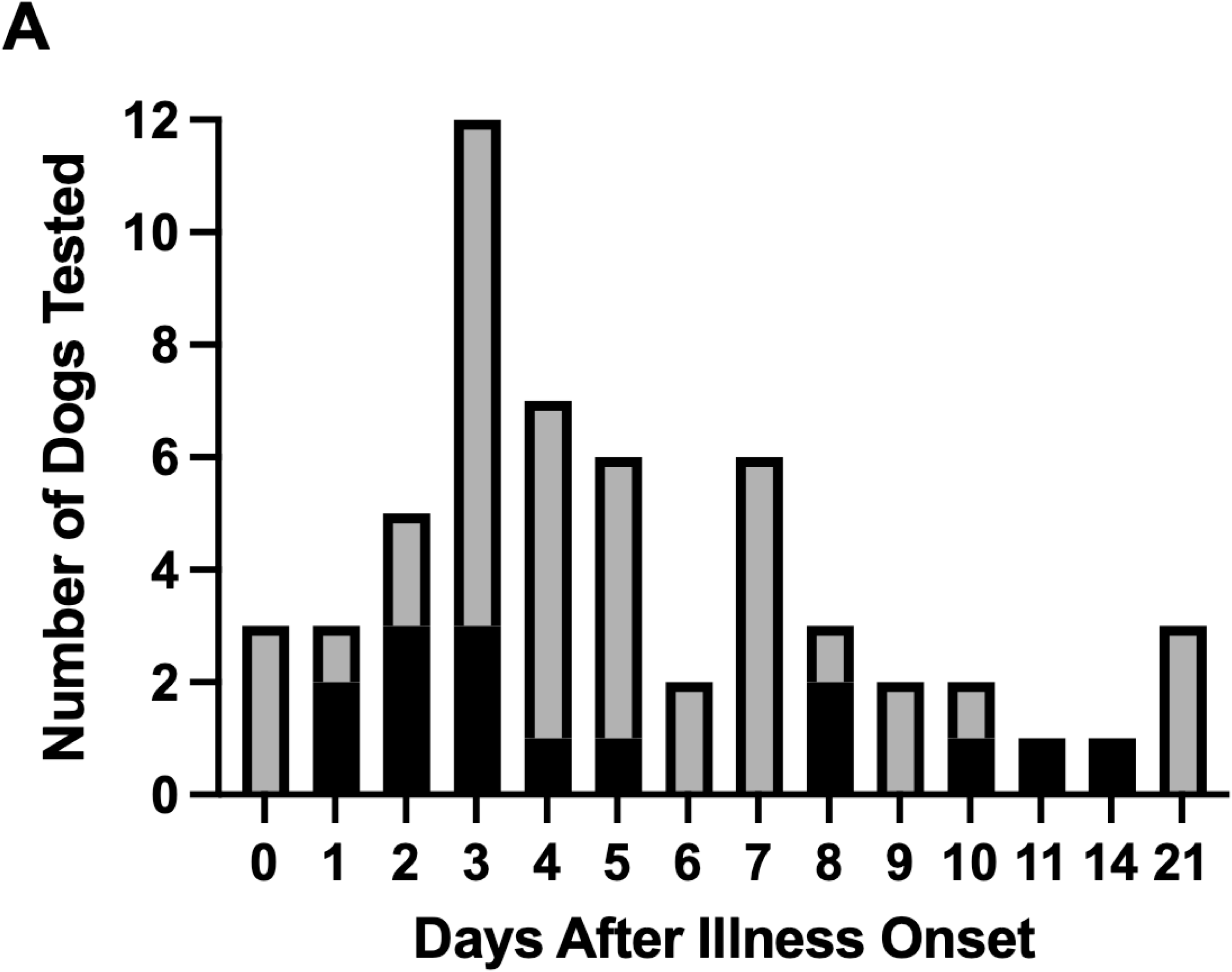
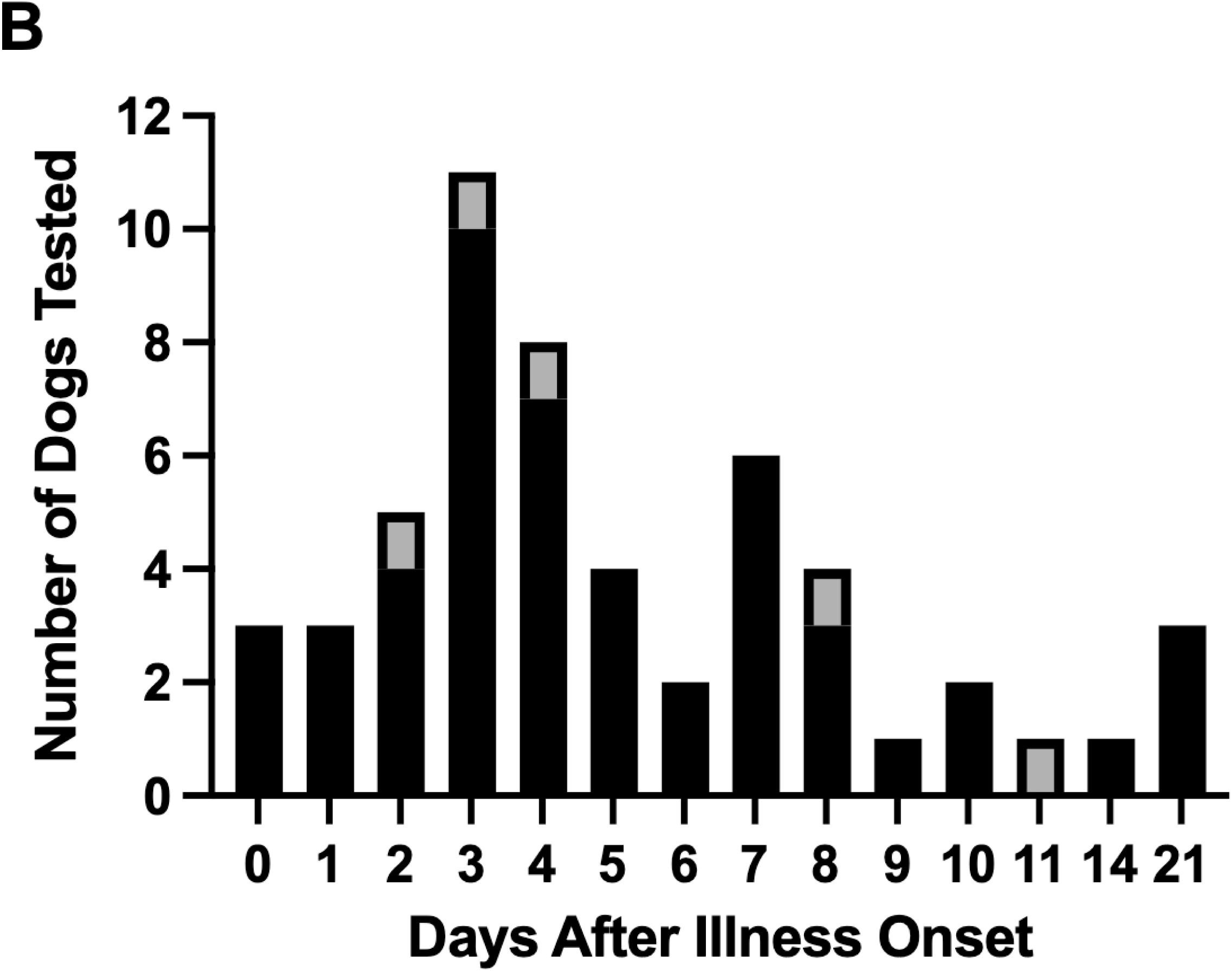
Results of *Leptospira LipL32* real-time PCR testing on blood (**A**) and urine (**B**) in 56 and 54 tested dogs with leptospirosis, respectively, plotted against the day of testing relative to illness onset. Black bars = number of dogs with positive results; grey bars = number with negative results.

### Serologic Testing using MAT

Serology using MAT was performed in 29/59 (49%) dogs, only one of which had recently received a leptospirosis vaccine (**Tables 3 and S5**). All seven dogs with negative MAT test results were tested from one to seven days after onset of illness. Of 20 dogs tested in the first week of illness, 9 had highest titers ≥ 1:6400; seven had negative titers. Nineteen (66%) of the 29 dogs tested had either a single highest titer ≥ 1:6400 (n = 8) or acute and convalescent testing (n = 11) with highest titers to serovar Canicola; lower titers were variably identified to other serovars tested. The acute to convalescent testing interval was 7 to 36 days (mean, 21 days; median, 23 days). Seroconversion was documented in six dogs. The remaining dogs had both high (> 1:6400) acute and convalescent titers, but acute and convalescent titers were either reported as the same or one dilution lower.

### Serologic Testing Using Point-of-Care (POC) Assays

Serum antibodies to *Leptospira* were detected in 20/27 (74%) dogs using the WITNESS Lepto Rapid Test (one dog had received a leptospirosis vaccine 10 days previously) and 8/9 (89%) dogs using the SNAP Lepto; one dog had both tests performed and was positive on both assays. Median (range) days of illness to test results are shown in **Table 3**. All except one dog with negative test results had been sick for ≤ 7 days; the single dog with a negative WITNESS test after 10 days of illness had a positive PCR result on blood, and illness began as intermittent vomiting and inappetence then transitioned to increased thirst and urination. Other serologic tests were not performed in this dog.

Using the case definition as the gold standard, the sensitivity of blood and urine PCR for *Leptospira* DNA was 27% and 91%, respectively. The sensitivity of POC assays (WITNESS Lepto Rapid Test and SNAP Lepto) for detection of antibodies to pathogenic leptospires was 74% and 89%, respectively. For initial MAT and when all serologic test types were combined, dogs with negative test results had a significantly shorter duration of illness than those with positive test results (median, 3 days versus 4 days, P = .02) (**Figure 5**).

**Figure 5.**
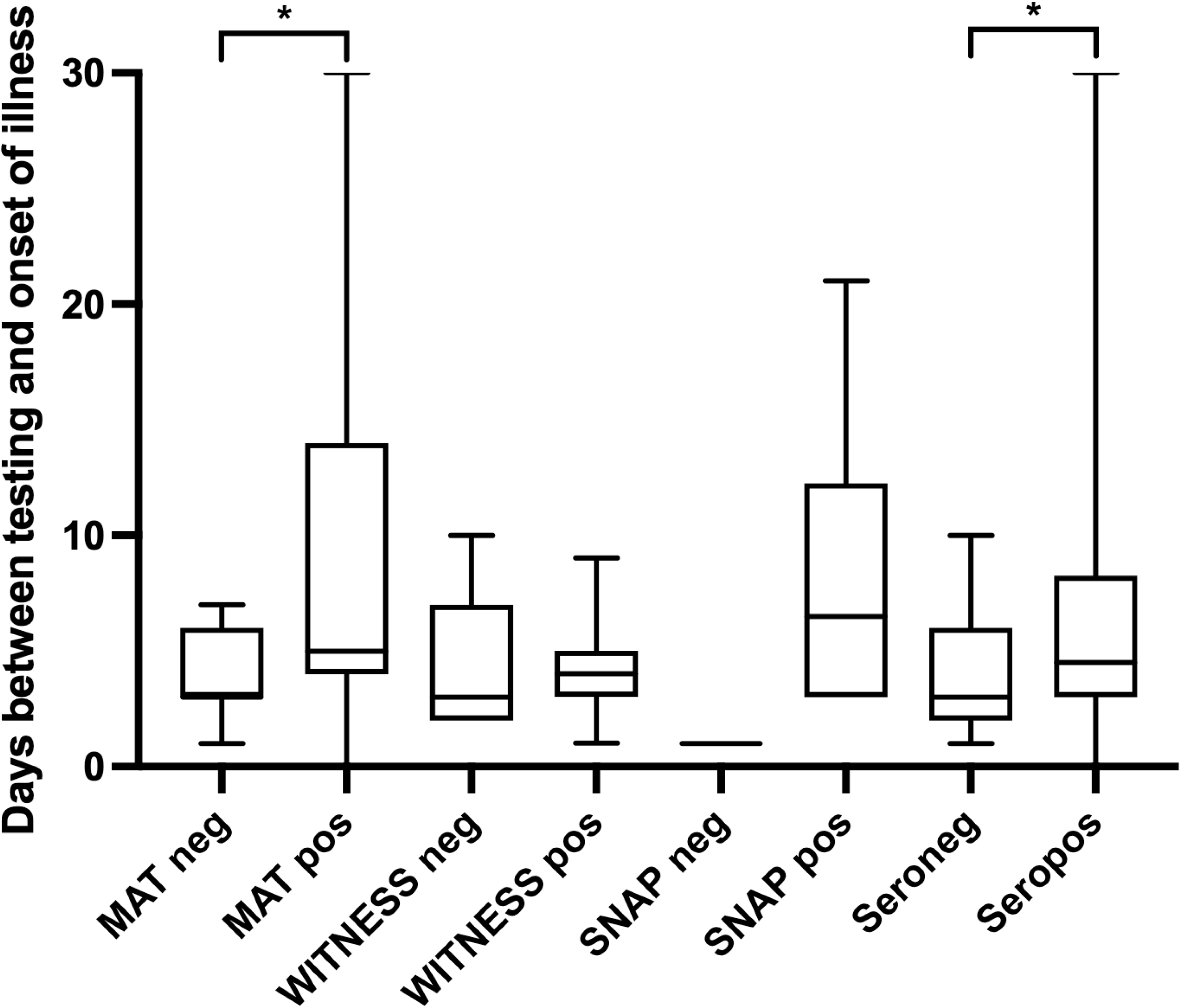
Box-and-whisker plots showing days from onset of illness to testing for microscopic agglutination test (MAT), the WITNESS Lepto Rapid test, the SNAP Lepto test, and all serologic test results in dogs with leptospirosis. neg=negative, pos=positive, * significant difference (.01 < P < .05). Whiskers represent the minimum to maximum values.

### Treatment and Outcome

Seventy-three percent (43/59) of dogs were hospitalized; 16 (27%) were treated as outpatients with oral doxycycline. Median duration of hospitalization was 4 (range, 1-15) days. Hospitalized dogs were treated with intravenous (IV) fluids and antibacterial drugs (IV ampicillin or ampicillin-sulbactam [n = 14], IV doxycycline [n = 14], oral doxycycline [n = 12], or a combination of ampicillin/sulbactam and doxycycline [n = 3]). Two dogs were treated with hemodialysis (nine treatments each). All dogs that lived to discharge (54/59 [92%]) were then treated with oral doxycycline for 7-30 (median 14, mean 15) days. For dogs that survived to follow-up, there was a progressive and significant reduction in median serum creatinine concentration from initial evaluation to discharge then discharge to follow-up (both P < 0.01) (**Table 4, Figure S2A and B**). The two dogs that were dialyzed had the highest initial serum creatinine concentrations (19.0 and 33.3 mg/dL); at last follow-up, these were 1.9 and 2.9 mg/dL, respectively. International Renal Interest Society (IRIS) AKI grades^44^ based on the last serum creatinine concentration available for dogs that lived and dogs that were euthanized are shown in **Figure S2C** (P < 0.001). The dog with IRIS Grade I AKI that was euthanized had suspected severe leptospiral pulmonary hemorrhage syndrome. The remaining four dogs that were euthanized had moderate to severe azotemia (IRIS AKI grades IV or V).

**Table 4.**
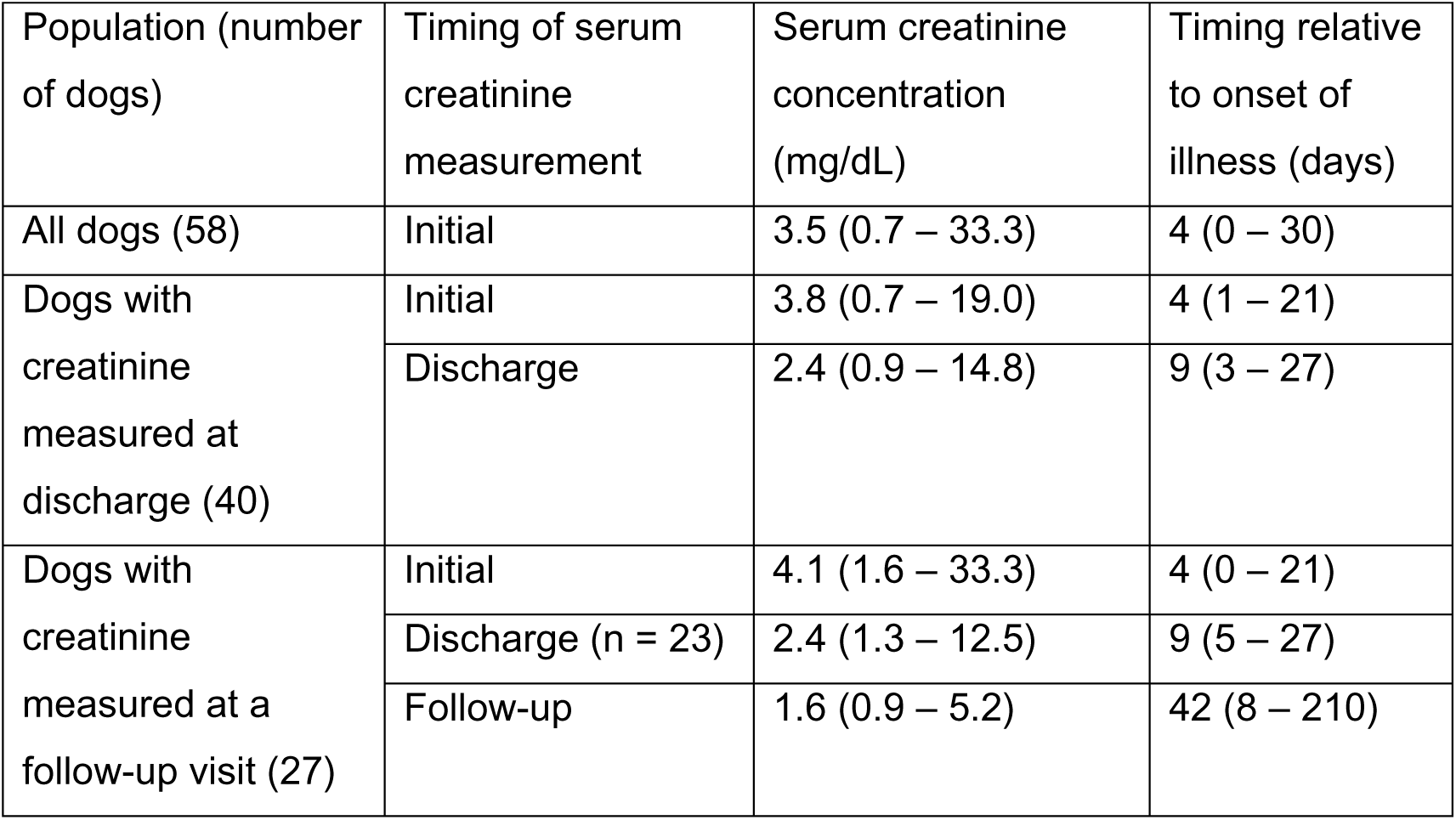
Median (range) of serum creatinine concentrations at initial evaluation, discharge, and follow-up relative to onset of illness in dogs diagnosed with leptospirosis during an outbreak in Los Angeles County.

## DISCUSSION

The results of this study offer significant advancements in our understanding of the epidemiology, diagnostic test performance, and treatment outcomes of leptospirosis in dogs, and shed light on approaches to reduction of a potential public health risk of dog ownership. The existence of an outbreak was supported by temporal and spatial clustering of cases. During the peak period, we found an association with ICF exposure. Although WGS analysis did not support dog-to-dog transmission, we could only obtain isolates during the post-peak period and only one of those isolates was from a dog with exposure to an ICF (boarding). However, many dogs had no history of ICF exposure or close contact with other dogs.

To our knowledge, this is the first study to report isolation and characterization of pathogenic leptospires from dogs from the United States since the early 1990s.^15,45^ Previous studies have relied on sequencing of PCR amplicons, which lacks discriminatory power and the ability to accurately identify the infecting serovar.^19^ All five dogs from which sequence information was available were infected with *L. interrogans* serovar Canicola ST34, but the infecting strains were distinct, suggesting multiple infection sources. Indeed, the LAD1-5 genomes represented 14.9% (143/960) of the total SNPs identified among the *L. interrogans* serovar Canicola genomes included in our phylogenetic analysis, which spanned 90 years and four continents. The two most similar genomes were from contiguous zip code regions 10 km east of the hotspot in Santa Monica; two other genomes collected 2 weeks apart from Santa Monica were less related. Infection of other dogs with serovar Canicola also seemed likely based on the highest titer to Canicola with a magnitude of ≥ 1:6400 in 17/22 dogs that tested positive with MAT. Detection of this serovar in sick dogs aligns with previous observations that dogs can act as either incidental or reservoir hosts for serovar Canicola,^4,46^ and supports the need for continued inclusion of serovar Canicola in vaccines for dogs in the United States. Whether serovar Canicola causes disease or persistent subclinical shedding in dogs might depend on pathogen virulence determinants and/or host factors. Through surveillance during the outbreak, the LA County DPH reported infection in six apparently healthy dogs.^25^ Of interest, *L. interrogans* serovar Canicola ST34 was recently identified in dogs from Mexico, including seven subclinically-infected dogs and two dogs with fatal leptospirosis.^4^

We identified temporal clustering of cases in July and August 2021. Although leptospirosis often occurs 1 to 3 months after heavy rainfall or flooding,^47^ rainfall in LA County from January-July 2021 was not above average.^48^ Another possible explanation for the timing of the outbreak is overcrowding of dog daycare facilities secondary to the rise in pet ownership during the COVID-19 pandemic, followed by return-to-work activity that was documented in Los Angeles in mid-2021.^49^ Concurrently, there was an outbreak of H3N2 influenza virus infection that involved over 1,300 dogs in west LA;^50^ such outbreaks typically follow importation of infected dogs from South Korea and China.^51^ A second possible pandemic-related explanation is increased rodent exposure, which occurred in large cities in the U.S. and other countries during the pandemic.^52,53^ In addition, the outbreak followed a massive (17 million gallon) offshore sewage spill into Santa Monica Bay, which occurred on July 11, 2021.^54^ Although it is unlikely this spill explained most cases based on timing and exposure histories, some dogs evaluated in the post-peak period had beach exposure in Santa Monica. Other urban wildlife species seemed unlikely to be a source based on a large study of specimens collected from striped skunks, raccoons, coyotes, fox squirrels, and Virginia opossums between 2015 to 2020.^31^ Using serology and WGS analysis, serovar Pomona appeared to be the predominant serovar in mesocarnivores; none of the species evaluated had significant seroreactivity to Canicola.

Historically, the risk of leptospirosis in urban dogs from Southern California has been considered low, so veterinarians were not routinely recommending leptospirosis vaccination. Evidence supports the efficacy and safety of four-serovar vaccines.^1^ Before four-serovar vaccines were available, dogs at-risk for leptospirosis were male, young adult, medium-to-large breed outdoor dogs in rural environments or that had recreational exposure to standing water.^1,55,56^ Since the availability of four-serovar vaccines, disease has predominantly been recognized in unvaccinated small breed dogs from urban regions.^1^ However, dogs in the outbreak reported here had risk factors that instead resembled those of the dog population with leptospirosis before the availability of four-serovar vaccines. It is possible that male dogs are more likely to exhibit behavior that increases their risk of contact with infected urine from rodents or other dogs. Among females, we found an association with intact status; estrogen can promote some other bacterial infections,^57^ but we were unable to find other studies of the effect of female gonadectomy on leptospirosis. The associations with young age and higher body weight were confounded by an interaction with boarding; this might have also impacted breed associations, as 6/7 dogs that were Siberian Huskies or Golden retrievers had exposure to ICFs. Breed associations might also reflect behavioral traits and/or breed-related immunodeficiency. Ultimately, the outbreak ended following educational efforts from the LAC DPH regarding the need for vaccination as well as attention to hygiene, rodent control, and pet population density in boarding facilities.

The results of diagnostic testing in this study supported the value of combining serologic test results and PCR to optimize diagnosis.^23^ Consistent with the results of previous studies,^58–60^ with the exception of the dog with a negative WITNESS test result at day 10 of illness, all dogs with negative serologic test results had been sick for ≤ 7 days. For dogs that failed to seroconvert, a rise in titer might have been missed because of a long median interval between titers (23 days rather than the recommended 7-14 days)^1^ or dogs might have been sick for longer than recognized by the owner. The latter seems likely based on the high titers in some dogs early in the course of illness. For some dogs that lacked serologic proof of infection, diagnosis was based on or a positive PCR result; however, given that healthy dogs can be leptospiruric, positive PCR results on urine alone require cautious interpretation.^1^ Because the WITNESS Lepto detects IgM, positive test results using this assay in dogs with consistent clinical signs are more likely to support a diagnosis of leptospirosis than assays that detect IgG (which are more likely to be positive from previous subclinical infection).^1^

The lower sensitivity of PCR testing on blood than on urine likely reflected the transient and early nature of leptospiremia.^1^ However, the frequent detection of leptospires in the urine of affected dogs was unexpected given that shedding by incidental hosts is often low-level and intermittent. The sensitivity of urine PCR might vary depending on host-pathogen interactions that influence the ability of the spirochete to persist in the renal tubules. The high prevalence of leptospiruria in this outbreak raises concern for dog-to-dog or dog-to-human transmission, although no evidence of human disease was documented by the LA County DPH.^25^

Although most dogs survived to discharge, renal function at discharge was variable and follow-up was limited. There was progressive recovery of renal function after discharge in some dogs. Dogs that were euthanized had severe azotemia or suspected leptospiral pulmonary hemorrhage syndrome. The high index of suspicion for the disease during the outbreak (both from owners and veterinarians) might have positively impacted outcomes and diagnostic test selection. Owner financial constraints likely also impacted outcome and test selection. Given that some recovered dogs remain infected with pathogenic leptospires despite antimicrobial treatment,^61,62^ and considering the particular ability of serovar Canicola to establish persistent infections in dogs, clinical recovery does not imply elimination of public health risk.

In summary, we were able to characterize an outbreak of leptospirosis in dogs, supporting the potential for dogs to act both as reservoir and incident hosts for serovar Canicola. Multiple independent infection sources seemed likely based on molecular and spatial epidemiologic analysis, which might have included exposure to other dogs in ICFs, rodents, or rodent urine. Further understanding would require studies such as characterization of isolates from rodent populations in the region. Early diagnosis using a combination of serology and PCR resulted in survival to discharge in over 90% of affected dogs. However, as this was a retrospective study that included only two referral clinics, other dogs involved in the outbreak might have died without a diagnosis, and dogs with negative initial PCR and serology results would not have been included in our study. Thus, conclusions cannot be made about the overall sensitivity and specificity of the diagnostic tests applied. Because analytical and clinical diagnostic test performance might be serovar-specific, findings might not apply to other regions or outbreak situations. Our findings support the need for widespread annual vaccination of all dogs to reduce the risk to dog and human health, a high index of suspicion for the disease in unvaccinated dogs, combined use of molecular and serologic tests to optimize diagnosis, and early treatment to optimize outcome.

## Abbreviations

AKI: acute kidney injury
ALP: alkaline phosphatase
ALT: alanine transaminase
ASEC: VCA Animal Specialty and Emergency Center, Los Angeles, CA
BUN: blood urea nitrogen
CI: confidence interval
Ct: cycle threshold
DFA: direct fluorescent antibody
DPH: Department of Public Health
HAN: Hornsby-Alt-Nally
LA: Los Angeles
ICF: indoor congregate facility
IV: intravenous
MAT: microscopic agglutination test
OR: odds ratio
PCR: polymerase chain reaction
POC: point-of-care
SNPs: single nucleotide polymorphisms
ST: sequence type
WGS: whole genome sequencing
WLA: VCA West Los Angeles Animal Hospital, Los Angeles, CA.

## ACKNOWLEDGMENT

No funding was received for this study. The authors acknowledge Marga Goris for her contributions posthumously.

## OFF-LABEL ANTIMICROBIAL DECLARATION

Antimicrobials were used off-label to treat dogs in this study as part of patient care.

## INSTITUTIONAL ANIMAL CARE AND USE COMMITTEE (IACUC) OR OTHER APPROVAL DECLARATION

No IACUC or other approval was needed as this was a retrospective study.

## HUMAN ETHICS APPROVAL DECLARATION

Human ethics approval was not needed for this study.

---

**Figure S1.** Spatio-temporal analysis of an outbreak of leptospirosis in dogs in Los Angeles County, including dogs evaluated at two specialty clinics (purple dots). **A.** Pre-peak. **B.** Peak. **C.** Post-peak. **D**. Intervals between case presentations over the course of the outbreak (range, 0 to 34 days, median = 2). Pre-peak, peak, and post-peak periods were selected based on frequency of case presentations at the two clinics (median and average for each period were 23.0 and 20.5 days, 1 and 1.8 days, and 3 and 4.6 days, respectively; pre-peak versus peak P < .01, peak versus post-peak P = .01, pre-peak versus post-peak P = .03). Asterisks indicate cases with a history of exposure to indoor congregate facilities (ICFs). **E** and **F.** Hot spot analysis for the peak and post-peak period, respectively. Open circles represent the number of cases with exposure to ICFs. The hotspot in Playa del Rey (90293) was associated with a high number of cases per household that did not have a history of exposure to ICFs.

**Figure S2. A.** Individual dog serum creatinine concentration over time in dogs diagnosed with leptospirosis that had more than one serum creatinine measurement, either at initial evaluation and discharge (40 dogs), at initial evaluation and a follow-up visit (27 dogs), or at initial evaluation, discharge, and a follow-up visit (23 dogs). **B.** Box-and-whisker plots showing serum creatinine concentration at time of initial evaluation, discharge, and final follow-up evaluation in 27 dogs with leptospirosis that had serial measurements performed. *Significant difference (P < 0.01). Whiskers represent the minimum to maximum values. **C.** International Renal Interest Society (IRIS) Acute Kidney Injury (AKI) grades based on last measured serum creatinine concentrations for dogs that lived to discharge (black columns) versus those that were euthanized (grey columns).

## REFERENCES

1. Sykes JE, Francey T, Schuller S, et al. 2023 Updated ACVIM consensus statement on leptospirosis in dogs. Journal of veterinary internal medicine / American College of Veterinary Internal Medicine 2023;37:1966–1982.

2. Sykes JE, Haake DA, Gamage CD, et al. A Global One Health Perspective on Leptospirosis in Humans and Animals. Journal of the American Veterinary Medical Association 2022;260:1589–1596.

3. Boey K, Shiokawa K, Rajeev S. Leptospira infection in rats: A literature review of global prevalence and distribution. PLoS neglected tropical diseases 2019;13:e0007499.

4. Carmona Gasca CA, Martinez Gonzalez S, Castillo Sanchez LO, et al. The Presence of a Virulent Clone of Leptospira interrogans Serovar Canicola in Confirmed Cases of Asymptomatic Dog Carriers in Mexico. Microorganisms 2024;12.

5. Schuller S, Moore GE, Sykes JE. Leptospirosis. In: Sykes JE, ed. Greene’s Infectious Diseases of the Dog and CatElsevier; 2023:802-823.

6. Goh SH, Khor KH, Radzi R, et al. Shedding and Genetic Diversity of Leptospira spp. From Urban Stray Dogs in Klang Valley, Malaysia. Top Companion Anim Med 2021;45:100562.

7. Koizumi N, Muto MM, Akachi S, et al. Molecular and serological investigation of Leptospira and leptospirosis in dogs in Japan. J Med Microbiol 2013;62:630–636.

8. Ward MP. Seasonality of canine leptospirosis in the United States and Canada and its association with rainfall. Prev Vet Med 2002;56:203–213.

9. Lee HS, Levine M, Guptill-Yoran C, et al. Regional and temporal variations of Leptospira seropositivity in dogs in the United States, 2000-2010. Journal of veterinary internal medicine / American College of Veterinary Internal Medicine 2014;28:779–788.

10. Lawson JH, Michna SW. Canicola fever in man and animals. Br Med J 1966;2:336–340.

11. Roczek A, Forster C, Raschel H, et al. Severe course of rat bite-associated Weil’s disease in a patient diagnosed with a new Leptospira-specific real-time quantitative LUX-PCR. J Med Microbiol 2008;57:658–663.

12. Nogueira DB, da Costa FTR, Bezerra CS, et al. Use of serological and molecular techniques for detection of Leptospira sp. carrier sheep under semiarid conditions and the importance of genital transmission route. Acta tropica 2020;207:105497.

13. Haake DA, Levett PN. Leptospirosis in Humans. Curr Top Microbiol Immunol 2015;387:65–97.

14. Ko AI, Goarant C, Picardeau M. Leptospira: the dawn of the molecular genetics era for an emerging zoonotic pathogen. Nat Rev Microbiol 2009;7:736–747.

15. Schmidt DR, Winn RE, Keefe TJ. Leptospirosis. Epidemiological features of a sporadic case. Arch Intern Med 1989;149:1878–1880.

16. Rosenberg BL. Canicola fever; review, with report of two new cases. Am J Med 1951;11:75–88.

17. Jorge S, Miotto BA, Kremer FS, et al. Complete genome sequence and in silico analysis of L. interrogans Canicola strain DU114: A virulent Brazilian isolate phylogenetically related to serovar Linhai. Genomics 2019;111:1651–1656.

18. Guzman DA, Diaz E, Saenz C, et al. Domestic dogs in indigenous Amazonian communities: Key players in Leptospira cycling and transmission? PLoS neglected tropical diseases 2024;18:e0011671.

19. Sykes JE, Gamage CD, Haake DA, et al. Understanding leptospirosis: application of state-of-the-art molecular typing tools with a One Health lens. Am J Vet Res 2022;83.

20. Santa Rosa CA, Sulzer CR, Yanaguita RM, et al. Leptospirosis in wildlife in Brazil: isolation of serovars canicola, pyrogenes and grippotyphosa. Int J Zoonoses 1980;7:40–43.

21. Bahaman AR, Ibrahim AL, Stallman ND, et al. The bacteriological prevalence of leptospiral infection in cattle and buffaloes in West Malaysia. Epidemiol Infect 1988;100:239–246.

22. Zacarias FG, Vasconcellos SA, Anzai EK, et al. Isolation of leptospira Serovars Canicola and Copenhageni from cattle urine in the state of ParanA, Brazil. Braz J Microbiol 2008;39:744–748.

23. Sykes JE, Reagan KL, Nally JE, et al. Role of Diagnostics in Epidemiology, Management, Surveillance, and Control of Leptospirosis. Pathogens 2022;11.

24. Los Angeles County Department of Public Health. http://www.publichealth.lacounty.gov/vet/Leptospirosis.htm. Last updated April 2022. In: 2022.

25. Los Angeles County Department of Public Health. http://www.publichealth.lacounty.gov/vet/Leptospirosis2021.htm. Last updated March 2022. Last accessed January 1, 2026,. In: 2022.

26. Hornsby RL, Alt DP, Nally JE. Isolation and propagation of leptospires at 37 degrees C directly from the mammalian host. Scientific reports 2020;10:9620.

27. Galloway RL, Hoffmaster AR. Optimization of LipL32 PCR assay for increased sensitivity in diagnosing leptospirosis. Diagn Microbiol Infect Dis 2015;82:199–200.

28. Stoddard RA, Gee JE, Wilkins PP, et al. Detection of pathogenic Leptospira spp. through TaqMan polymerase chain reaction targeting the LipL32 gene. Diagn Microbiol Infect Dis 2009;64:247–255.

29. Hamond C, LeCount K, Browne AS, et al. Concurrent colonization of rodent kidneys with multiple species and serogroups of pathogenic Leptospira. Appl Environ Microbiol 2023;89:e0120423.

30. Hartskeerl R, Smits H, Korver H, et al. International Course on Laboratory Methods for the Diagnosis of Leptospirosis. Course ManualRoyal Tropical Institute, Amsterdam, The Netherlands; 2006.

31. Helman SK, Tokuyama AFN, Mummah RO, et al. Pathogenic Leptospira are widespread in the urban wildlife of southern California. Scientific reports 2023;13:14368.

32. Stone NE, McDonough RF, Hamond C, et al. DNA Capture and Enrichment: A Culture-Independent Approach for Characterizing the Genomic Diversity of Pathogenic Leptospira Species. Microorganisms 2023;11.

33. Wood DE, Lu J, Langmead B. Improved metagenomic analysis with Kraken 2. Genome Biol 2019;20:257.

34. Bankevich A, Nurk S, Antipov D, et al. SPAdes: a new genome assembly algorithm and its applications to single-cell sequencing. J Comput Biol 2012;19:455–477.

35. Insitut Pasteur. BIGS-db Pasteur. https://bigsdb.pasteur.fr/cgi-bin/bigsdb/bigsdb.pl?db=pubmlst_leptospira_seqdef&page=sequenceQuery. Last accessed January 6, 2026. In.

36. Li H. Minimap2: pairwise alignment for nucleotide sequences. Bioinformatics 2018;34:3094–3100.

37. McKenna A, Hanna M, Banks E, et al. The Genome Analysis Toolkit: a MapReduce framework for analyzing next-generation DNA sequencing data. Genome Res 2010;20:1297–1303.

38. Delcher AL, Phillippy A, Carlton J, et al. Fast algorithms for large-scale genome alignment and comparison. Nucleic Acids Res 2002;30:2478–2483.

39. Sahl JW, Lemmer D, Travis J, et al. NASP: an accurate, rapid method for the identification of SNPs in WGS datasets that supports flexible input and output formats. Microb Genom 2016;2:e000074.

40. Nguyen LT, Schmidt HA, von Haeseler A, et al. IQ-TREE: a fast and effective stochastic algorithm for estimating maximum-likelihood phylogenies. Mol Biol Evol 2015;32:268–274.

41. Kalyaanamoorthy S, Minh BQ, Wong TKF, et al. ModelFinder: fast model selection for accurate phylogenetic estimates. Nat Methods 2017;14:587–589.

42. Li H, Handsaker B, Wysoker A, et al. The Sequence Alignment/Map format and SAMtools. Bioinformatics 2009;25:2078–2079.

43. International Leptospirosis Proficiency Testing Scheme. 2026. https://leptosociety.org/proficiency_testing/. Last accessed January 7, 2026. In.

44. International Renal Interest Society (IRIS). https://www.iris-kidney.com/iris-guidelines-1. Last accessed January 6, 2026. In: 2025.

45. Anderson JF, Miller DA, Post JE, et al. Isolation of Leptospira interrogans serovar grippotyphosa from the skin of a dog. Journal of the American Veterinary Medical Association 1993;203:1550–1551.

46. Miotto BA, Guilloux AGA, Tozzi BF, et al. Prospective study of canine leptospirosis in shelter and stray dog populations: Identification of chronic carriers and different Leptospira species infecting dogs. PloS one 2018;13:e0200384.

47. Chadsuthi S, Chalvet-Monfray K, Wiratsudakul A, et al. The effects of flooding and weather conditions on leptospirosis transmission in Thailand. Scientific reports 2021;11:1486.

48. Los Angeles Almanac. Historical Monthly Rainfall by Season. Santa Monica, California (Santa Monica Airport). https://www.laalmanac.com/weather/we139aa.php. Last accessed January 2026. In: 2021.

49. CBRE Group, Inc. LA’s office re-entry slows despite signs of strength in demand. https://cbreemail.com/cv/8b499a22a8c210320968bc32b4ba91850efc846e. Last accessed January 2026. In: 2021.

50. Los Angeles County Department of Public Health. Canine Influenza. http://publichealth.lacounty.gov/vet/influenzacanine.htm. Last updated March 2022. Last accessed January 2026. 2022.

51. Voorhees IEH, Dalziel BD, Glaser A, et al. Multiple Incursions and Recurrent Epidemic Fade-Out of H3N2 Canine Influenza A Virus in the United States. J Virol 2018;92.

52. Murray MH, Byers KA, Buckley J, et al. “I don’t feel safe sitting in my own yard”: Chicago resident experiences with urban rats during a COVID-19 stay-at-home order. BMC Public Health 2021;21:1008.

53. Orkin. The Rat Pack: Former Top Five Cities Remain Leaders on Orkin’s 2021 Rattiest Cities List. https://www.orkin.com/press-room/orkin-top-rattiest-cities-2021. Last accessed January 2026. 2021.

54. California Water Environment Association. https://www.cwea.org/news/las-hyperion-experiences-overwhelming-flood-of-debris-clogging-headworks/. August 13, 2021. Last accessed January 7, 2026. In.

55. Ward MP, Glickman LT, Guptill LE. Prevalence of and risk factors for leptospirosis among dogs in the United States and Canada: 677 cases (1970-1998). Journal of the American Veterinary Medical Association 2002;220:53–58.

56. Adin CA, Cowgill LD. Treatment and outcome of dogs with leptospirosis: 36 cases (1990-1998). Journal of the American Veterinary Medical Association 2000;216:371–375.

57. Hong L, Liang H, Man W, et al. Estrogen and bacterial infection. Front Immunol 2025;16:1556683.

58. Lizer J, Grahlmann M, Hapke H, et al. Evaluation of a rapid IgM detection test for diagnosis of acute leptospirosis in dogs. Vet Rec 2017;180:517.

59. Lizer J, Velineni S, Weber A, et al. Evaluation of 3 Serological Tests for Early Detection Of Leptospira-specific Antibodies in Experimentally Infected Dogs. Journal of veterinary internal medicine / American College of Veterinary Internal Medicine 2018;32:201–207.

60. Gloor CI, Schweighauser A, Francey T, et al. Diagnostic value of two commercial chromatographic “patient-side” tests in the diagnosis of acute canine leptospirosis. The Journal of small animal practice 2017.

61. Juvet F, Schuller S, O’Neill EJ, et al. Urinary shedding of spirochaetes in a dog with acute leptospirosis despite treatment. Vet Rec 2011;168:564.

62. Hetrick K, Harkin KR, Peddireddi L, et al. Evaluation by polymerase chain reaction assay of persistent shedding of pathogenic leptospires in the urine of dogs with leptospirosis. Journal of veterinary internal medicine / American College of Veterinary Internal Medicine 2022;36:133–140.

