## Supplemental Table 1 to 4 for "Clinical and molecular characterization of an outbreak of leptospirosis in dogs from Los Angeles County, California, USA, 2021"

**Table S1**. Accession numbers for sequences obtained from five dogs infected with *Leptospira interrogans* serovar Canicola as detected using culture or genome enrichment from urine. All sequences were deposited under BioProject number PRJNA1377681.

| **Dog identification** | **Biosample accession** | **SRA accession** | **Assembly accession** |
| --- | --- | --- | --- |
| LAD1 | SAMN53827873 | SRR36381965 | JBSYGR00000000 |
| LAD2 | SAMN53827874 | SRR36381964 | NA |
| LAD3 | SAMN53827875 | SRR36381963 | JBSYGQ00000000 |
| LAD4 | SAMN53827876 | SRR36381962 | JBSYGP00000000 |
| LAD5 | SAMN53827877 | SRR36381961 | JBSYGO00000000 |

NA, not applicable.

**Table S2**. Sampling dates, zip codes, and exposure histories for five dogs infected with *Leptospira interrogans* serovar Canicola as detected using culture or genome enrichment from urine. Zip codes are spatially illustrated in Figure 3.

| **Dog identification** | **Specimen collection date** | **Method of isolation** | **Zip code of residence** | **Region** | **Exposure history** |
| --- | --- | --- | --- | --- | --- |
| LAD1 | 10/4/2021 | Culture | 90046 | West Hollywood | Neighbors’ dogs |
| LAD2 | 10/10/2021 | Genome enrichment | 90036 | West Hollywood | Dog park |
| LAD3 | 10/13/2021 | Culture | 90066 | Mar Vista | Boarding or dog park |
| LAD4 | 11/17/2021 | Culture | 90405 | Santa Monica/Venice | Unknown |
| LAD5 | 11/22/2021 | Culture | 90405 | Santa Monica/Venice | Boarding |

**Table S3**. Possible sources of exposure to *Leptospira* spp. in 59 dogs diagnosed with leptospirosis at two specialty clinics during an outbreak in Los Angeles County.

| **Possible Source*** | **Period** | | | **Total (n)** |
| --- | --- | --- | --- | --- |
|  | **Pre-peak (n)** | **Peak (n)** | **Post-peak (n)** |  |
| **Indoor Congregate Facilities** |  |  |  |  |
| Dog daycare | 1 | 19 | 3 | **22** |
| Long-term boarding | 0 | 2 | 3 | **5** |
| Grooming or training | 0 | 2 | 0 | **2** |
| Dog shows | 0 | 1 | 0 | **1** |
| TOTAL INDOOR | 1 | 24 | 6 | **31** |
| **Outdoor or Unknown** |  |  |  |  |
| Dog parks | 0 | 1 | 5 | **6** |
| Neighborhood walks or hikes | 2 | 1 | 3 | **6** |
| Beaches | 0 | 0 | 2 | **2** |
| Travel out of state† | 1 | 0 | 1 | **2** |
| Rodent ingestion | 0 | 1 | 0 | **1** |
| No known exposure | 0 | 3 | 2 | **5** |
| Unknown | 1 | 4 | 3 | **8** |
| TOTAL OTHER | 4 | 10 | 16 | **30** |
| **TOTAL** | **5** | **34** | **22** | **61** |

n, number of dogs.
*one dog had exposure to both dog parks and boarding, and another beaches and dog parks.
†Arizona and the Caribbean.

**Table S4**. Results of initial serum biochemistry testing for renal and hepatic function in dogs diagnosed with leptospirosis at two specialty clinics during an outbreak in Los Angeles County. Results are expressed as median (range).

| Biochemical findings | Creat  (mg/dL) | ALP  (IU/L) | ALT  (IU/L) | TBili  (mg/dL) | Duration of illness (days) |
| --- | --- | --- | --- | --- | --- |
| Concurrent renal and hepatic involvement (n = 6) | 3.6 (1.5 – 4.1) | 197 (58 – 1137) | 286 (32 – 1427) | 1.9 (0.2 – 11.8) | 3 – 6 |
| Isolated renal involvement (n = 29) | 3.6 (1.6 – 19.0) | 49 (13 – 152) | 31 (16 – 92) | 0.2 (<0.1-0.3) | 0 – 21 |
| Reference Interval | 0.4-1.3 | 5-131 | 12-118 | <0.1-0.3 |  |

Creat, creatinine; ALP, alkaline phosphatase; ALT, alanine transaminase; TBili, total bilirubin.
