## Supplemental Figure 1 for "Clinical and molecular characterization of an outbreak of leptospirosis in dogs from Los Angeles County, California, USA, 2021"

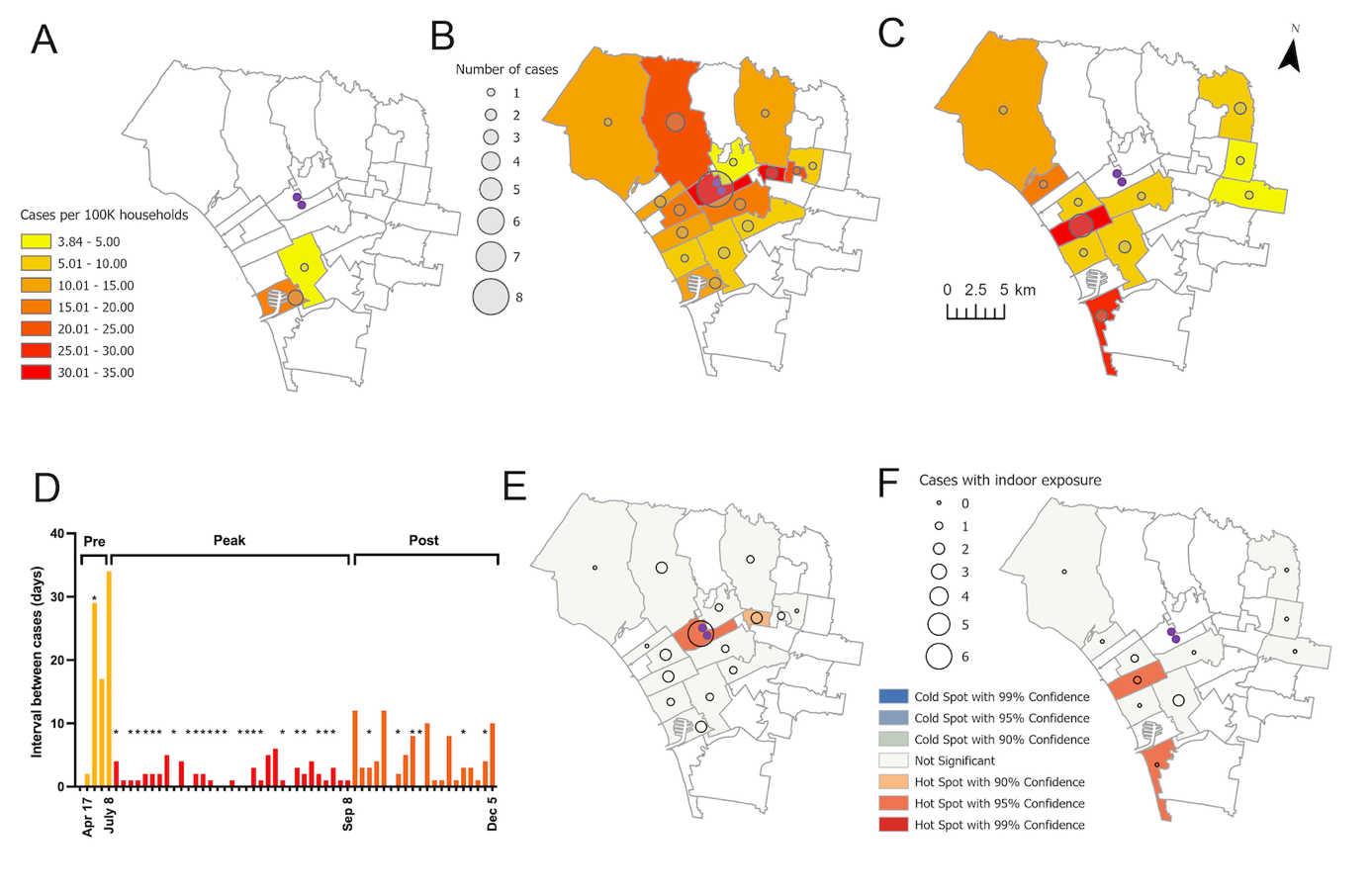

**Figure S1.** Spatio-temporal analysis of an outbreak of leptospirosis in dogs in Los Angeles County, including dogs evaluated at two specialty clinics (purple dots). **A.** Pre-peak. **B.** Peak. **C.** Post-peak. **D**. Intervals between case presentations over the course of the outbreak (range, 0 to 34 days, median = 2). Pre-peak, peak, and post-peak periods were selected based on frequency of case presentations at the two clinics (median and average for each period were 23.0 and 20.5 days, 1 and 1.8 days, and 3 and 4.6 days, respectively; pre-peak versus peak P < .01, peak versus post-peak P = .01, pre-peak versus post-peak P = .03). Asterisks indicate cases with a history of exposure to indoor congregate facilities (ICFs). **E** and **F.** Hot spot analysis for the peak and post-peak period, respectively. Open circles represent the number of cases with exposure to ICFs. The hotspot in Playa del Rey (90293) was associated with a high number of cases per household that did not have a history of exposure to ICFs.
