## Supplemental Figure 2 for "Clinical and molecular characterization of an outbreak of leptospirosis in dogs from Los Angeles County, California, USA, 2021"

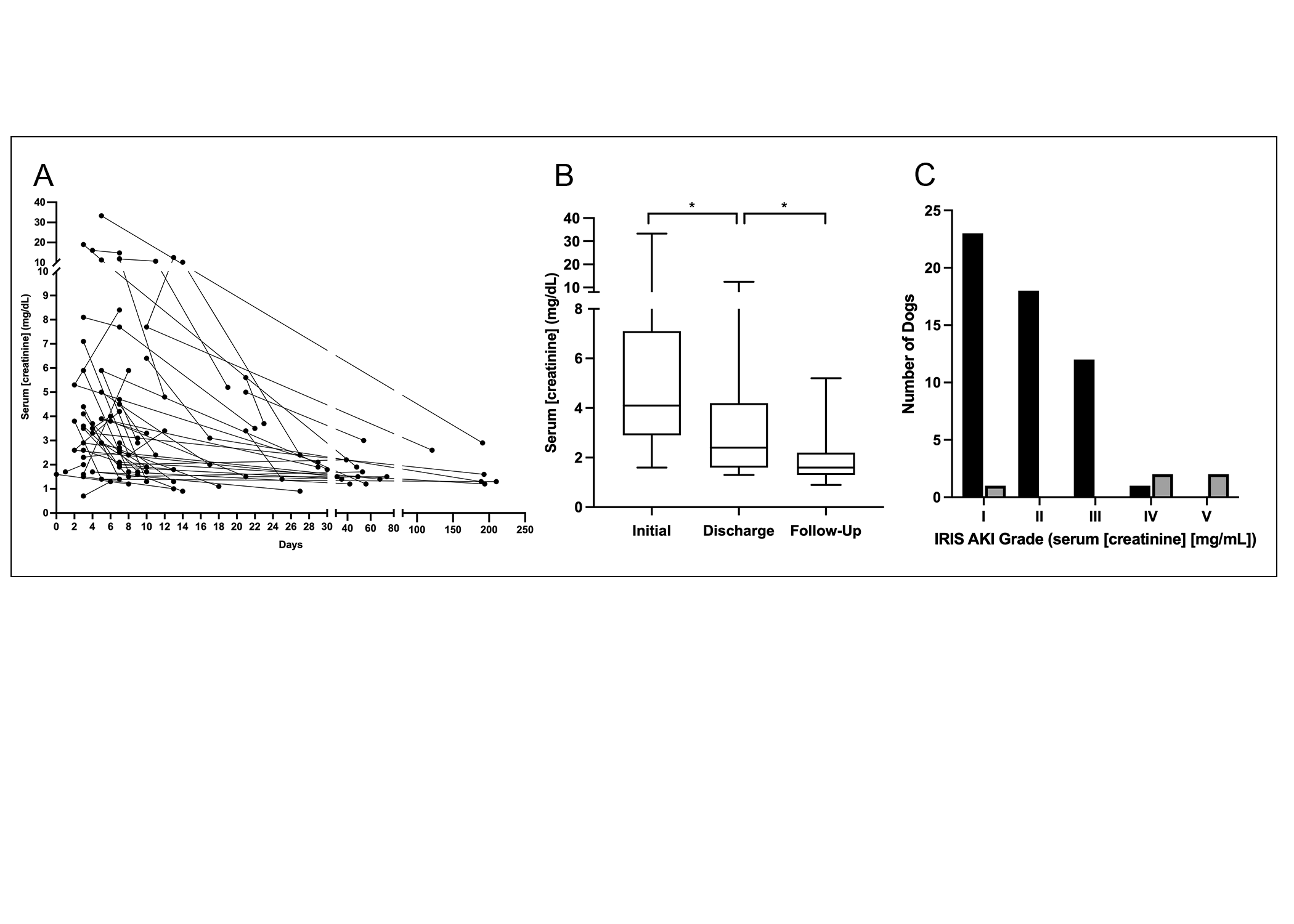

**Figure S2. A.** Individual dog serum creatinine concentration over time in dogs diagnosed with leptospirosis that had more than one serum creatinine measurement, either at initial evaluation and discharge (40 dogs), at initial evaluation and a follow-up visit (27 dogs), or at initial evaluation, discharge, and a follow-up visit (23 dogs). **B.** Box-and-whisker plots showing serum creatinine concentration at time of initial evaluation, discharge, and final follow-up evaluation in 27 dogs with leptospirosis that had serial measurements performed. *Significant difference (P < 0.01). Whiskers represent the minimum to maximum values. **C.** International Renal Interest Society (IRIS) Acute Kidney Injury (AKI) grades based on last measured serum creatinine concentrations for dogs that lived to discharge (black columns) versus those that were euthanized (grey columns).
